# Adaptive Reinforcement Learning is causally supported by Anterior Cingulate Cortex and Striatum

**DOI:** 10.1101/2024.06.12.598723

**Authors:** Robert Louis Treuting, Kianoush Banaie Boroujeni, Charles Grimes Gerrity, Adam Neumann, Paul Tiesinga, Thilo Womelsdorf

## Abstract

Learning the reward structure of complex environments can be achieved using reinforcement learning processes augmented with cognitive strategies. Among these strategies is the adjustment of the exploration-exploitation trade-off to increase exploration when behavior gets stuck and increase exploitation when reward contingencies remain stable. Here we tested how the anterior cingulate cortex (ACC) and the striatum causally support adaptive cognitive strategies augmenting reinforcement learning. We electrically microstimulated the ACC or the striatum in nonhuman primates at the time they chose multidimensional objects to learn about their reward values, while varying target feature uncertainty and the motivational saliency of the chosen objects. We found that stimulation of the ACC and the striatum affected adaptive strategies and reinforcement learning when target feature uncertainty was high, but in opposite ways. ACC-stimulation impaired learning and sustaining correct responses, while striatum-stimulation on average improved learning from rewarding outcomes. Behavioral modeling showed that stimulation affected the same mechanisms but in opposite ways. ACC stimulation impaired the monitoring of outcome uncertainty for adapting exploration-exploitation and reduced the ability to lower prediction errors during learning, while striatum stimulation enhanced the monitoring of outcome uncertainty and prediction-error based updating of object values. Stimulation did not alter the use of working memory or attentional filtering as alternative learning strategy. These opposing behavioral stimulation effects were associated with the ACC having populations of neurons that fired stronger during choices that were more uncertain, had a lower value, and that tracked error history, while striatum neurons more likely encoded higher values and more certain choices. In summary, microstimulation during object choices suggest that ACC and the striatum causally adapt exploration-exploitation levels to guide exploration towards reward-relevant objects during periods of uncertainty.

**Short Summary:** The anterior cingulate cortex (ACC) and the striatum are core nodes of a network supporting reinforcement learning, but how they augment reinforcement learning with adaptive cognitive strategies has remained unresolved. This study uses electrical microstimulation to show that ACC and the striatum have causal roles adjusting the exploration-exploitation balance and optimize reinforcement learning in multidimensional environments, while not altering attentional filtering or working memory strategies.

## Introduction

Reinforcement learning (RL) reduces the uncertainty about the value of options. This uncertainty reduction can be inefficient in volatile environments and in multidimensional environments with many possible reward-relevant features as is typical for the real world. For these volatile and multidimensional environments, reinforcement learning can be augmented with adaptive strategies and extended to incorporate cognitive strategies ^1,2^. Previous studies found that in multidimensional environments, subjects’ learning progress is better predicted with adaptive strategies that allowed for more exploration and dynamic learning rates when value uncertainty increases, or that allows working memory to guide decision making when the expected values of options are ambiguous or low ^3–7^. These computational insights suggest that adaptive learning strategies might be supported by brain systems tracking the uncertainty of outcomes, such as the anterior cingulate cortex (ACC) ^8,9^, and that integrate reward information across different learning systems, such as the striatum ^10^. Here, we set out to test how the ACC and the striatum causally contribute to adaptive RL by transiently microstimulating these areas while subjects learned to reduce reward uncertainty in a multidimensional learning task.

Studies testing the causal contribution of the ACC to learning have shown that disrupting the ACC impairs the ability to choose among objects in volatile environments when object values vary probabilistically and are uncertain ^11–13^, but impairments are also apparent in learning and set shifting paradigms that require shifting choices between objects that have otherwise fixed reward associations ^14–17^. For all these paradigms, ACC disruption does not result in persevering with the same non-optimal choices, but rather in a difficulty sustaining attending to and choosing objects once they are rewarded. It has been suggested that one reason for this pattern of deficits after ACC disruption is an impaired integration of reward information over time to estimate the value of currently available objects ^18,19^ and to guide the next choice to the option with the highest predicted value ^12,20^. It has remained unresolved, however, whether the ACC’s role in tracking reward history is used to adaptively change RL during periods of uncertainty.

One possibility to adaptively improve RL is to use cognitive strategies such as working memory (WM) of recently rewarded objects ^4,6,21^. Both the ACC and the striatum have been implicated in WM processes. Striatal lesions impair not only reward learning but also WM ^22^, while a role of the ACC in WM can be inferred from a disruption study that found the ACC mediates choosing counterfactual stimuli that are not available as options for a current choice but which would be the best valued alternative option in future trials based on their past values ^12^. The maintenance of object values for unavailable options reflects a WM process that computational studies have shown to augment reinforcement learning processes when subjects learn values in environments with multiple multidimensional objects^17,23^ or learn complex sensori-motor associations ^3,6,21,24,25^. While these studies suggest that WM processes support learning, it is unknown whether the ACC or striatum causally contribute to a learning strategy that combines WM and reinforcement learning.

Another versatile way to improve RL learning in volatile and multidimensional environments is to optimize the exploration-exploitation balance given the currently experienced levels of outcome uncertainty ^25,26^. Both the ACC and the striatum may support this mechanism. The ACC has been implicated in guiding exploration during periods of uncertainty in learning experiments ^9,25,27^ and prolonged ACC disruptions impairs exploratory fixational sampling of objects ^17^, suggesting that the ACC may be able to dynamically adjust the exploration-exploitation balance during learning. In the striatum, neuronal firing tracks during learning the certainty of value-based choice ^28^ and choice outcomes ^29^, while dopaminergic tone modulates the degree of exploration to exploitation^30^. Direct microstimulation of the anterior striatum increases the value of an object that is processed during the stimulation event and has been shown to increase exploitation of that object ^31^, while object-specific stimulation had no effect in another study ^32^. Whether these acute stimulation effects could support adaptive reinforcement learning in multidimensional environments that require object specific credit assignment processes remains unresolved.

To address these questions and determine possible casual contributions of the ACC and the striatum to learning in multidimensional environments, we devised a task that increased the demands on exploration vs exploitation of objects by varying the number of object features subjects needed to evaluate during learning. In addition, we varied the motivational saliency of choosing objects by varying the gains and losses for correct and erroneous choices. We found that microstimulation affected learning particularly when feature uncertainty was high. Microstimulation in the ACC disrupted learning and in the striatum it improved learning. Computational modeling ^23^ suggests that stimulation affected not only standard reinforcement learning mechanisms but also the exploration-exploitation tradeoff, inducing more exploration when stimulating the ACC, and inducing more efficient exploitation of values when stimulating the striatum. The pattern of results suggest that the ACC and striatum are causally important for enabling adaptative reinforcement learning in complex, multidimensional environments.

## Results

We applied in each of two monkeys electrical microstimulation to the fundus of the anterior cingulate cortex (area 24c) and to the striatum (head of the caudate) (48 sessions per subject, 24 sessions per area). Microstimulation was delivered during at least the first twelve trials and continued until subjects learned the new feature-reward rule. A brief 0.3 s stimulation pattern was initiated when monkeys fixated an object for longer than 0.3 s, which indicated choosing that object, and receive either token rewards or losses for that choice (**Figure 1A**). In a given block, stimulation was administered either when the rewarded target object was chosen (*StimReinf+* condition), or when the non-rewarded object was chosen (*StimReinf-* condition) (**Figure 1A**). Monkeys learned which object was rewarded through trial-and-error. In each trial three objects were shown. Subjects explored them with fixational eye movements until they chose an object by sustaining fixation of an object for 0.7 s. Choices were followed by visual and auditory feedback and by the addition of tokens for correct responses or the subtraction of already attained tokens for incorrect responses. Tokens were green coin symbols in a token bar presented at the top of the screen. Accumulating eight tokens resulted in fluid reward. The rewarded target feature remained constant for a block of 30-50 trials before it switched to a new feature that was not shown before (**Figure 1B**). Blocks differed pseudo-randomly in their motivational context (+2 or +3 tokens gained for correct choices; loss of –1 or –3 tokens for errors; token outcomes were deterministically provided for choices with an outcome probability of 1), and their learning difficulty. Difficulty levels corresponded to the size of the feature space, i.e. the number of visual features that distinguished objects, increasing from features of one, two, to three feature dimensions (1D, 2D, and 3D learning conditions) (**Figure 1C**). Monkeys F / I completed on average 33.5 / 36 blocks during ACC stimulation sessions and 32.0 / 35.8 blocks during striatum stimulation sessions.

**Figure 1.**
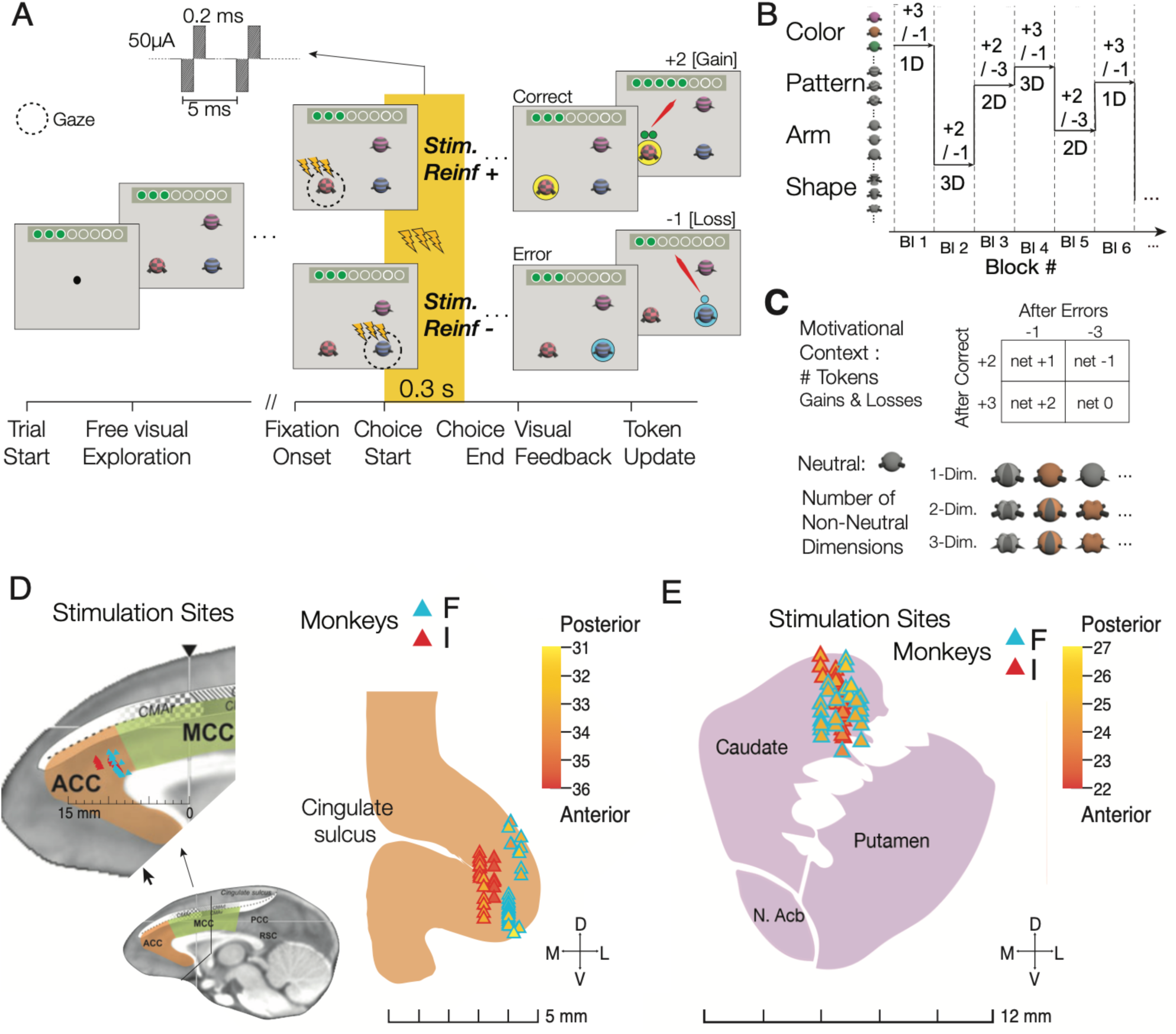
Task paradigm and stimulation locations. (**A**) On each trial monkeys freely explored three stimuli until they fixated on one stimulus for longer than 0.3 s which indicated they committed choosing that stimulus. Electrical (or sham) stimulation was delivered 0.3-0.6 s after the onset of the fixation of the chosen stimulus. Stimulation was triggered either when the positive reinforced (rewarded) object was chosen (*StimReinf+)* or when a negative reinforced stimulus (resulting in loosing tokens) was chosen (*StimReinf-).* Stimulation was only delivered during learning new feature-reward associations, i.e. prior to reaching a 70% performance criterion. 0.75 s after the choice-fixation onset a halo behind the chosen object provided visual feedback (yellow/blue for correct/error choices), followed 0.5 s later by auditory feedback and the reveal of tokens gained or lost. (**B**) The task switched the rewarded feature (y-axis) in blocks of 30-50 trials. (**C**) Blocks randomly varied the motivational context from gaining +2 or +3 for choosing the object with the rewarded feature, and the loss of 1 or 3 tokens otherwise. Blocks also varied randomly whether features from one, two, or three feature dimensions of the objects changed from trial to trial (1D, 2D, and 3D conditions). (**D**) Stimulation locations in the ACC for monkeys F / I (blue / red) in the reference frame of Procyk et al, 2016 (left) and shown on a coronal view (right). Coloring indicates the anterior-posterior distance to the interaural line. (**E**) Stimulation locations in the striatum (head of the caudate nucleus) for monkeys F / I (blue / red).

### Microstimulation sites within a connected cingulate-striatal circuit

The anatomical locations for microstimulation in the ACC (**Figure 1D**) focused on the cognitive subfield of the ACC that has prominent connectivity to the lateral prefrontal cortex, the anterior striatum, and executive function networks ^33–35^. Neural activity at this ACC site codes for values of covertly attended stimuli ^36^, values of choice options ^37,38^, and prediction errors of visual features ^29,39^. The prediction error responses suggest this ACC site tracks previous trials’ outcomes which could inform triggering a behavioral change of choice strategies when outcomes are unfavorable or unexpected ^29,39–41^. The anatomical locations for microstimulation in the anterior striatum (**Figure 1E**) focused on the head of the caudate nucleus, which has neurons with activity tracking the expected value of visual objects during learning ^28,42^ and is anatomically connected to the ACC ^43–45^, with neuronal spiking activity synchronizing with spiking activity in the ACC during attention, choice, and outcome periods ^35,46^.

### Microstimulation affects learning in the multidimensional environment

Without microstimulation (sham condition) monkeys successfully learned feature-reward rules at >70% correct in the 1D/2D/3D conditions of the ACC sessions after an average of 4.0/11.6/20.2 trials (monkey F) and 4.0/11.3/15.02 trials (monkey I) and in the striatum sessions after an average of 4.0/11.3/19.7 trials (monkey F) and 3.9/10.8/18.8 trials (monkey I) (**Figure 2**). Microstimulation systematically affected learning of both subjects in both areas. The average performance curves of each of the monkeys showed that in the difficult 3D task condition the learning criterion was reached later when the ACC was microstimulated (**Figure 2A**), and learning criterion was reached earlier when the striatum was microstimulated (**Figure 2B**), while there was no consistent modulation for the easier 1D and 2D conditions. The specific microstimulation effects on learning in the multidimensional (3D) condition were similarly found when using different thresholds for defining the trials-to-criterion and confirmed statistically. ACC microstimulation overall impaired learning dependent on dimensionality (2-way ANOVA: stim. condition x dimensionality interaction, Fstat = 3.06, p = 0.0159). Impaired learning was evident in each subject in the 3D condition in blocks with microstimulation in correct trials (*StimReinf+*) compared to sham, but not with stimulation during erroneous choices (*StimReinf-*) (**Figure 2C**) (3D condition *StimReinf+* vs sham, monkey F: Fstat = 4.15, p = 0.0448; monkey I: Fstat = 5.94, p = 0.0166; *StimReinf-* vs sham: monkey F: Fstat = 1.89, p = 0.1733; monkey I: Fstat = 0.43, p = 0.5121). Monkeys F and I needed on average 3.05 and 8.19 trials longer to reach criterion performance in the *StimReinf+* than in the sham condition (trials-to-criterion in the *StimReinf+* / *StimReinf-* / sham condition for monkey F: 23.26 (SE: 2.54) / 19.30 (SE: 2.27) / 20.21 (SE: 1.70); for monkey I: 23.22 (SE: 2.71) / 17.29 (SE: 1.95) / 15.03 (SE: 1.40)). While monkey F appeared to have slower learning in the 2D condition with *StimReinf-* stimulation (**Figure 2C**), this did not reach significance (FDR-corrected pairwise t-test for *StimReinf-* vs sham, monkey F: p = 0.1224).

**Figure 2.**
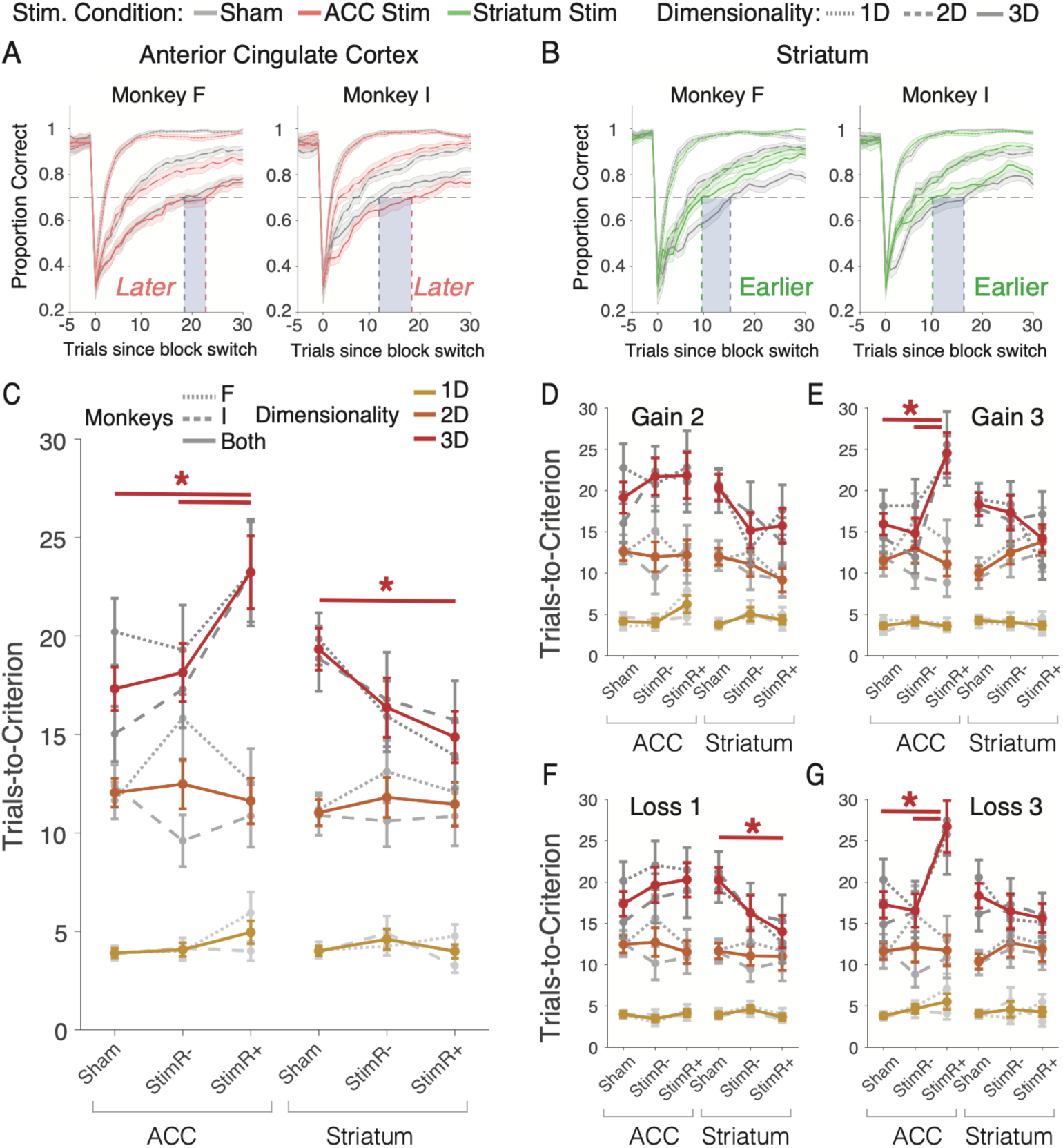
Stimulation in anterior cingulate cortex impairs and in striatum improves learning in multidimensional (3D) conditions. (**A**,**B**) Average Learning curves for blocks with stimulation (*StimReinf+* and *StimReinf-* combined) and sham in the ACC (A) and the striatum (B) for each monkey in the 1D, 2D, and 3D condition. Vertical grey lines denote the trial at which performance reached criterion performance (first trial with 70% correct over subsequent 12 trials) in the 3D condition, which was *later* with stimulation vs sham in the ACC and *earlier* with stimulation vs sham in the striatum. (**C**) Average number of trials to reach 70% performance criterion (y-axis) for the 1D/2D/3D dimensionality conditions for the sham, *StimReinf+*, *StimReinf-* stimulation conditions in the ACC (left) and the striatum (right). Colored lines reflect both monkeys, grey lines show individual monkeys. Horizontal bars mark slower learning in *StimReinf+* than sham condition for high dimensionality blocks. (**D-G**) Same format as C for 1D/2D/3D blocks where correct trials led to gains of +2 tokens (*D*), +3 tokens (*E*), and when erroneous choices led to a loss of 1 token (*F*) and 3 tokens (*G*). Error bars are SEM. Asterisks (*) indicate FDR-corrected p < 0.05 using t-tests with both monkeys pooled.

In the striatum, microstimulation affected behavior also as a function of dimensionality, but with an opposite sign relative to the ACC (**Figure 2C**). Striatal microstimulation overall improved learning dependent on dimensionality (2-way ANOVA: stimulation condition x dimensionality interaction, Fstat = 2.78, p = 0.0256). Learning was improved in the 3D condition in blocks with stimulation in correct trials (*StimReinf+*) compared to sham, but not with stimulation during erroneous choices (*StimReinf-*) in monkey F (**Figure 2C**) (3D condition *StimReinf+* vs sham, monkey F: Fstat = 3.96, p = 0.0494; *StimReinf-* vs sham: monkey F: Fstat = 3.22, p = 0.0756), while monkey I showed a qualitatively similar result that did not reach statistical significance (*see* below; 3D condition *StimReinf+* vs sham, monkey I: Fstat = 1.57, p = 0.2137; *StimReinf-* vs sham: monkey I: Fstat = 0.17, p = 0.6787). Monkeys F and I were on average 5.93 and 3.12 trials faster to reach criterion performance in the *StimReinf+* than in the sham condition (trials-to-criterion in the *StimReinf+* / *StimReinf-* / sham condition for monkey F: 13.92 (SE: 1.68) / 15.92 (SE: 1.80) / 19.85 (SE: 1.33); for monkey I: 15.73 (SE: 1.99) / 16.78 (SE: 2.29) / 18.85 (SE: 1.65).

We next tested whether microstimulation affected learning differently in blocks with high versus low gains (gain of 3 tokens in G3-L1 and G3-L3 blocks versus 2 tokens in G2-L1 and G2-L3 blocks,) or losses (blocks with loss of 1 versus 3 tokens, i.e. G2-L1 and G3-L1 versus G2-L3 and G3-L3). In the ACC, there was no 2-way interaction of microstimulation with either the gain or loss factor, but there was a 3-way interaction with dimensionality and gains, signifying overall slower learning in the ACC in the 3D condition with high versus low gains (**Figure 2D,E**) (3-way ANOVA, overall: Fstat = 2.41, p = 0.0476). Post-hoc t-tests showed that this effect reflected slower learning in blocks with larger gains primarily in monkey I, and less but qualitatively similar in monkey F (**Figure 2E**) (trials-to-criterion with gains of 3 token in the *StimReinf+* / sham condition for monkey I: 25.5 / 14.3, p = 0.0053; for monkey F: 23.6 / 18.1, p = 0.3304). There was no overall 3-way interaction of stimulation condition x dimensionality x loss (**Figure 2F,G**) (3-way ANOVA, overall: Fstat = 1.25, p = 0.2801). However, finer visual inspections showed that in the ACC learning slowed with *StimReinf+* in the 3D conditions in blocks with high loss (–3 tokens) (**Figure 2G**) (FDR-corrected t-test, p = 0.0099). This effect in blocks with higher loss was qualitatively similar in both monkeys (trials-to-criterion in the *StimReinf+* / sham conditions for monkey F: 25.8 / 20.3, p = 0.4003; for monkey I: 27.4 / 14.9, p = 0.0099).

In the striatum, there was no 2-way or 3-way interaction of microstimulation with either varying gains or losses and dimensionality (**Figure 2D-G**). Visual inspection showed the only improvement was evident in the low (–1 tokens) loss condition in 3D condition with microstimulation on correct trials (*StimReinf+*), where overall (monkeys combined) learning was faster, but this effect was not significant in either animal (FDR-corrected t-test, overall: p = 0.0496; trials-to-criterion in the *StimReinf+* / sham conditions for monkey F: 12.6 / 9.1, p = 0.0756; for monkey I: 15.3 / 21.2, p = 0.3977) (**Figure 2F**).

The impairment with ACC microstimulation and the improvement with striatum microstimulation in the 3D condition specifically affected the learning of new feature-reward associations. Microstimulation (*1*) did not change the overall likelihood that the learning criterion was reached within a block (**Figure S1A**), (*2*) did not affect the plateau performance once learning was completed (**Figure S1B**), (*3*) had no overall effect on choice reaction times during learning or when learning was completed (**Figure S1C,D**), (*4*) did not have a systematic influence on the number of objects subjects fixated (sampled) prior to choosing an object before or after reaching criterion performance (**Figure S1E,F**), and (*5*) did not systematically alter the duration of object fixations prior to making a choice (**Figure S2A,B** and **Supplemental Results**). There were only selective conditions when the task was difficult (3D conditions) and the gains and losses were low in which stimulation of the striatum resulted in shorter choice reaction times or reduced number of object fixations with reduced viewing durations (**Figure S1C-F, Figure, S2B**).

The stimulation effects on learning might vary over time within and across experimental sessions. We found that stimulation slowed learning in ACC and improved learning in the striatum in the 3D condition across twenty-four sessions and from earlier to later blocks during sessions in monkey I, while monkey F showed more prominent slowing of learning (ACC) in earlier sessions and in the first two thirds of experimental sessions (**Figure S2C-J**). One source of varying efficacy of the stimulation effects could be how strong the stimulation pulses modulated neural activity at a particular stimulation site. We extracted the post-stimulation multiunit activity peaks in the stimulation and sham conditions for individual sessions and correlated them with the behavioral stimulation effects on learning. However, we found no correlation between the strength of the stimulation effects on neural activity and behavioral learning in the ACC (**Figure S3A-D**), or the striatum (**Figure S3E-H**).

### Subjects similarly contribute effective stimulation sessions to the average stimulation effects

Another source of variability of stimulation effects is the anatomical site of stimulation. Prior studies have reported heterogeneous effects of stimulation in the ACC and in the striatum on approach/avoidance behavior with anatomically nearby sites having sometimes opposite stimulation effects on how stimuli are approached ^47,48^. We evaluated the anatomical heterogeneity of stimulation effects by quantifying the effect of microstimulation on learning in the 3D relative to the 1D condition for individual experimental sessions. This analysis revealed some heterogeneity of stimulation effects but also confirmed an overall prevalence of sessions showing impaired learning with ACC microstimulation (**Figure 3A**) and improved learning with striatum microstimulation (**Figure 3B**). We calculated the number of effective sessions as those sessions for which the behavioral stimulation effect was statistically significant and found that each subject contributed a similar proportion of effective sessions to the main effects. In the ACC, microstimulation impaired learning in 7 (29%) and 10 (42%) sessions in monkey F and I, respectively, while learning was improved in fewer sessions (3 and 2 in monkey F and I, respectively; improved vs impaired sessions in ACC for both monkeys combined: p = 0.0036; for monkeys F / I: p =0.1551 / p = 0.0077, Chi-square tests) (**Figure 3C**). In the striatum, effective sessions were also consistent with an overall learning improvement. Most effective sessions in the striatum showed improved learning (8 and 9 in monkey F and I, 33% and 38%, respectively) while fewer sessions (n=2, monkey F) or a similar number of sessions (n=9, monkey I) showed impaired learning (**Figure 3D**) (Chi-square test, improved vs impaired sessions in monkeys combined: p = 0.0363, for monkey F: p =0.0330, for monkey I: p = 0.3502). We confirmed that the impairment in the effective stimulation sessions in the ACC was selectively evident in the 3D condition (**Figure S4A-E**). Similarly, sessions with improved learning in the striatum showed the improvement specifically in the 3D task condition (**Figure S4F-J**). In separate analyses we tested whether sessions with a significant improvement or impairment of learning were clustered anatomically in the ACC or the striatum, but did not find anatomical clustering of these functional effects.

**Figure 3.**
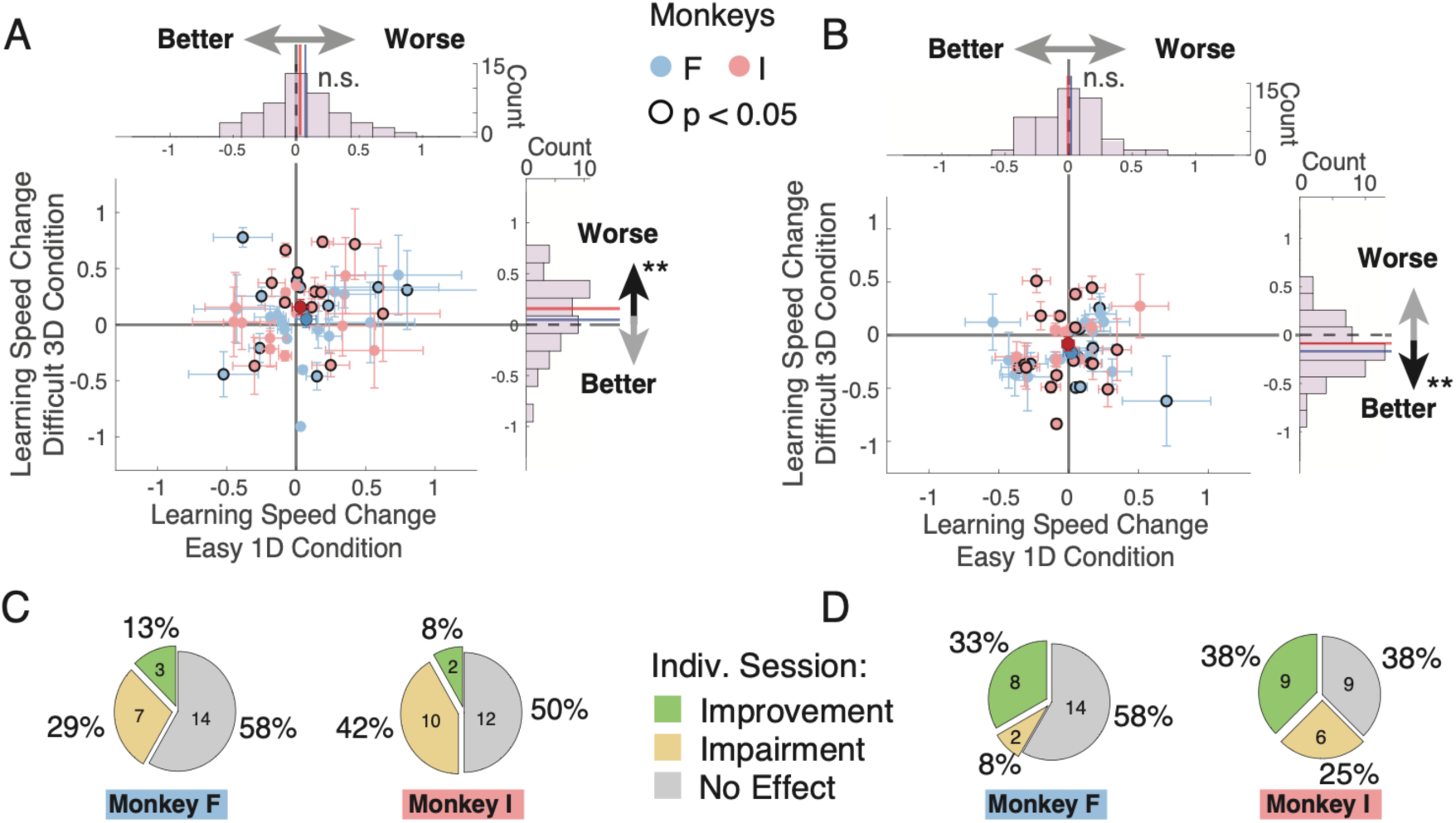
The majority of stimulation sessions in the ACC impairs learning and in the striatum improves learning. (**A**) The scatterplot shows for each experimental session in the ACC the difference in learning speed (trial-to-criterion) with stimulation vs sham in the 3D condition (*y-axis*) and the 1D condition (*x-axis*). Black outlined markers denote sessions with individually significant difference of stim vs. sham. Blue /Red data are from subject F/I. Errors bars are STD. Solid point. is mean. Histogram summarized the effects for 1D (upper) and 3D (rightmost) condition. Values <0 shows slower learning with stimulation than sham. (**B**) Pie charts summarizing the proportion of sessions in which stimulation improved (green) or impaired (yellow) learning for each subject. (**C,D**) Same format as (*A*,*C*) for experiments in the striatum.

### Stimulation affects positive credit assignment

The results so far show that robust stimulation effects on learning particularly with microstimulation in rewarded trials (*StimReinf+* condition) (**Figure 2C-G**, **S4A-J**), which suggest that stimulation modulates the positive credit assignment process that associates visual feature with the positive reward expectations, and less with the assignment of negative credit after error trials, i.e. recognizing which features were incorrectly chosen and should be avoided. We tested this suggestion by focusing on the 3D condition on trials in a block prior to reaching the learning criterion and quantifying how subjects used outcomes for the trial-by-trial adjustment of performance following correct trials (i.e. after gaining tokens) compared to error trials (i.e. after losing tokens). The analysis was done separately for correct and error trials in the *StimReinf+* and the *StimReinf-* condition. In the ACC, microstimulation in the *StimReinf+* condition overall impaired performance around trials with gains, while there was no consistent effect after losses (main effect of stimulation (β2) for gains: t-stat = –2.7409, p = 0.0061; for losses: t-stat = –1.594, p = 0.1109) (**Figure 4A,B**). Similarly, for the *StimReinf-* condition an impairment was visible as a trend around gains, but there was no effect around losses (**Figure 4C,D**) (main effect of stimulation (β2) for gains: t-stat = –1.91, p = 0.0561, t-stat = –0.8764; for losses: p = 0.3807). These trial-by-trial effects around gains and losses of *StimReinf+* and *StimReinf-* in the ACC were evident in each subject (**Figure S5**). Restricting the analysis to the 17 effective sessions confirmed that stimulation during rewarded (*StimReinf+)* and non-rewarded (*StimReinf-*) trials impaired performance prominently around gains, but not reliably after losses (**Figure S4K-N**).

**Figure 4.**
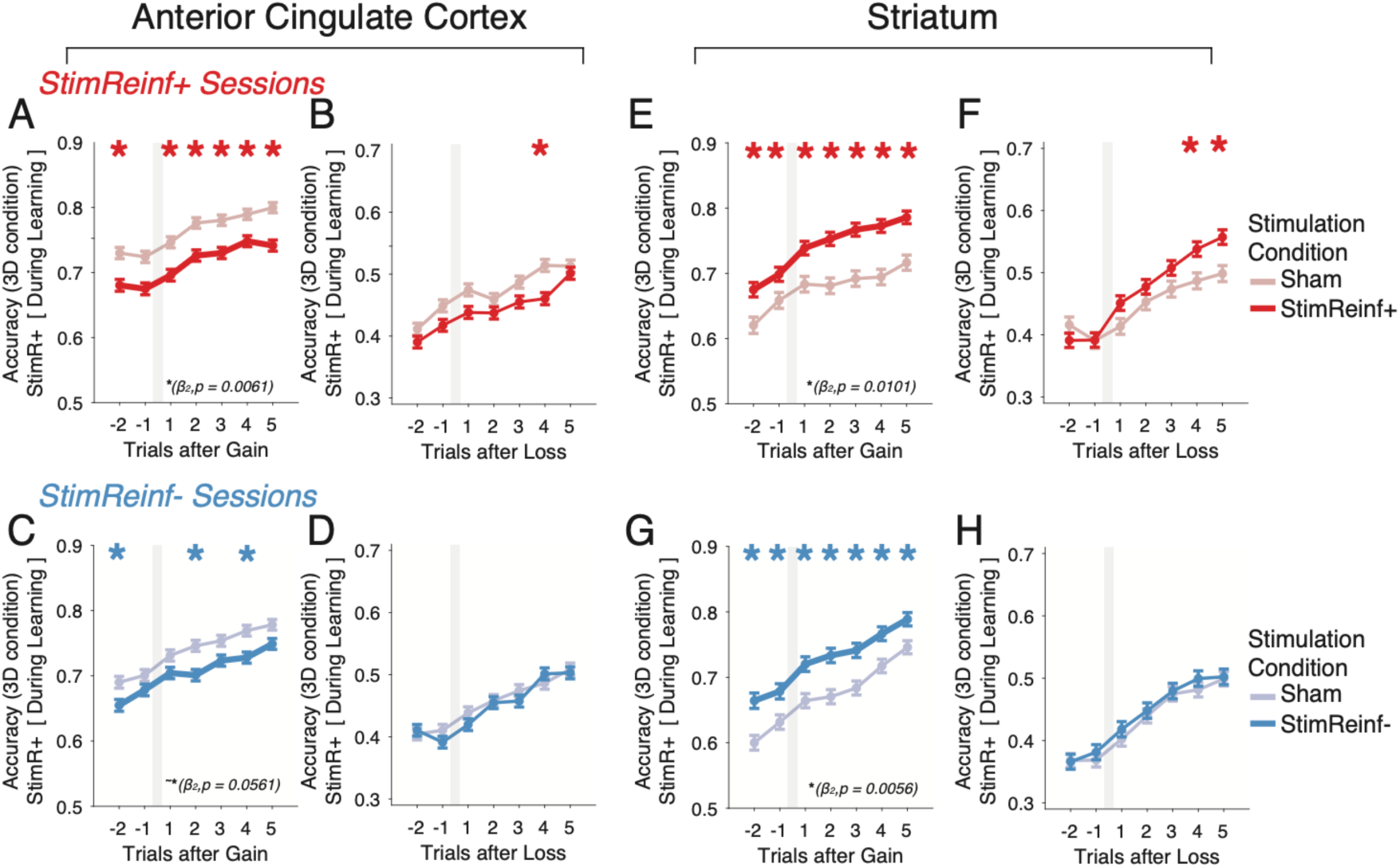
Stimulation modulates behavior following gains in the 3D condition. (**A**) Average accuracy (*y-axis*) for trials before and after a correct trial that gained subjects tokens for blocks with stimulation in the ACC on correct trials (*StimReinf+*) and sham in the 3D condition for trials until the mean trials criterion in the ACC / *StimReinf+* condition. *’s denote FDR-corrected p<0.05 (permutation test) significant differences of stim vs sham. The p-value at bottom refers to a significant coefficient using the Wald test after stimulation and sham conditions were fitted to a logistic regression model, produced from a generalized linear model with a binomial distribution and logit link function (see Methods). (**B**) Same as (*A)* for accuracy before and after incorrect trials that lost tokens. (**C,D**). Same format as (*A*,*B*) for experimental sessions with *StimReinf-* stimulation and sham. (**E-H**). Same format as (*A-D*) for experimental sessions in the striatum with *StimReinf+* stimulation (*E*,*F*), and with *StimReinf-* stimulation (*G*,*H*).

A similar result pattern was evident in the striatum but with the opposite sign relative to the ACC. Microstimulation in the *StimReinf+* and in the *StimReinf-* condition improved trial-by-trial adjustment following gains (**Figure 4E,G**), but around losses, there was only an effect when restricting the analysis to the 17 sessions with a stimulation induced learning improvement (for *StimReinf+*: main effect of stimulation (β2) for gains: t-stat = 2.5716, p = 0.0101; for losses: t-stat = 0.7338, p = 0.4631; for *StimReinf-*: main effect of stimulation (β2) for gains: t-stat = 2.7683, p = 0.0056; for losses: t-stat = 0.5640, p = 0.5727) (**Figure 4F,H; Figure S4O-R**). The result pattern was similar in both subjects (**Figure S5**). Taken together, these findings suggest that microstimulation in the ACC interferes, and in the striatum facilitates most prominently with positive credit assignment, and that similar effects for negative credit assignment become evident only in the striatum when considering sessions with the strongest effect sizes.

The trial-by-trial effects of microstimulation on performance during learning of feature-reward associations raises the question how fast the brief 0.3 s electrical stimulation pulses affected behavior. Statistically reliable behavioral effects of microstimulation in the 3D condition emerged after the sixth stimulation trial in the ACC in the *StimReinf+* condition (**Figure S6A,B**), while in the striatum performance improvements became evident already after the second (*StimReinf+*) and first (*StimReinf-*) stimulation trial (**Figure S6C,D**).

### Microstimulation in ACC disrupts sustaining correct responses

Lesions studies suggest that an intact ACC is important for sustaining correct responses ^11,14^, and that the striatum supports preventing persevering non-rewarded choices ^49^. To test whether microstimulation affected these behavioral patterns we quantified how stimulation changed the likelihood of correct choices in trials after committing one error (after an ‘E’), after a single correct trial (‘EC’), after two correct trials (‘ECC’), etc. This EC_n_ analysis showed that in the ACC microstimulation reduced performance after sequences of three correct choices (‘ECCC’) in the *StimReinf+* condition, consistent with a disruption of sustaining rewarded choices (FDR-correct p-values < 0.05, permutation test; **Figure S6E**). There was no EC_n_ effects with microstimulation in the striatum (**Figure S6G,H**). There were no differences of stimulation compared to sham conditions with regard to sequences of erroneous choices (CE_n_ analysis).

### Microstimulation modulates reinforcement learning and adaptive exploration

Microstimulation impaired versus improved learning after stimulation of the ACC versus the striatum. At the cognitive-behavioral level this dissociation could either indicate that the ACC and the striatum implement different learning mechanisms, or that they modulate the same learning mechanisms but in an opposite way. To evaluate these alternatives we fit the trial-by-trial performance of the subjects with models that account for choices of subjects with either one, two, or three computationally distinct learning mechanisms that previous studies have shown to be important for learning in environments with multidimensional objects ^2,4,23,50^. For this analysis we focused on the 3D condition and the effective sessions where stimulation impaired learning in the ACC (**Fig. 3B**) and improved learning in the striatum (**Fig. 3D**).

We evaluated as first mechanism a Rescorla Wagner RL model augmented with a selective forgetting (SF) parameter. This ‘*SF model*’ used a parameter (ω^*selectiveDecay*^) to decay the values of features from non-chosen objects akin to an attentional attenuation of non-chosen stimuli, which is a mechanism that speeds up learning among multidimensional options ^23,29,50^ (**Figure 5A**). As a second mechanism we evaluated a RL model with adaptive exploration (AE). This ‘*AE model*’ tracked prediction errors over trials in a separate error-trace variable, which it weighted with an error-trace weight (α^ErrorTrace^) that increased exploration over exploitation when erroneous choices accumulated, and reduced exploration when errors were rare ^23,25,51^. The mechanism reflects meta-learning insight about the level of experienced outcome uncertainty (in the form of prediction errors), increasing exploratory choices when uncertainty increases. As third mechanism we evaluated working memory (WM), which supports learning by uploading an object when its choice led to reward, decaying its representation with parameter ω^+,^, and biases choosing objects that have the strongest WM representation (see methods). WM can support fast learning as it rapidly updates objects after an outcome without needing to compute prediction errors for slowly updating value expectations ^3,4,6^. We evaluated each of the three model mechanisms and their combinations (**Figure 5B**). For models combining the RL models (i.e. the SF or AE model) with the WM module we used a weighting variable (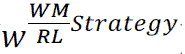), that determined for each trial how strong the RL component contributed to choices relative to the objects represented in WM (with values lower than 0.5 reflecting a larger influence of RL) (*see* Methods).

**Figure 5.**
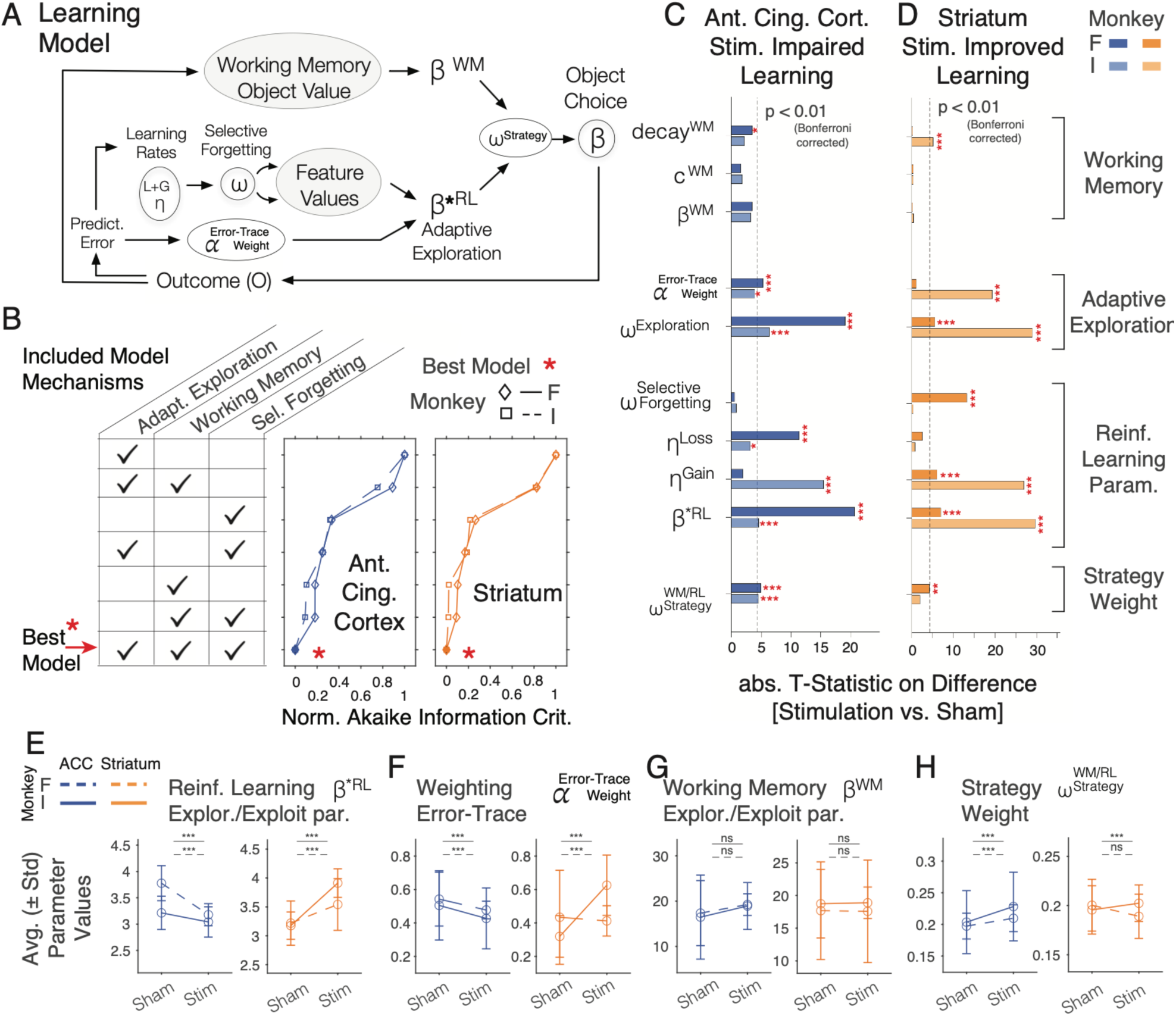
A triple-strategy model accounts for learning at high (3D) feature uncertainty. (**A**) Overview of the triple-strategy model combining adaptive exploration / selective forgetting / working memory (AE-SF-WM). The model learns to make choices of objects by a combination of (i) working memory (WM) and by using expected feature values of an RL module, which uses (ii) an adaptive mechanism to adjust exploration during learning and (iii) a selective forgetting of values from non-chosen features. (**B**) The AE-SF-WM model best fit choices in the effective high (3D) feature-uncertainty condition across sessions and stimulation / sham conditions in the ACC (blue) and striatum (orange) for both monkeys (solid and dashed line). The lowest normalized Akaike Information Criterion corresponds to the best model. (**C,D**) The effect size (absolute value of t-statistics) and Bonferroni corrected p-values for the difference of parameter values of the model fit separately to the sham and stimulation conditions in the 3D condition in ACC (*C*) and in the striatum (*D*). The parameters are defined in Methods. (**E-H**) Change in fitting parameters between stimulation and sham condition for ACC (blue) and striatum (orange) for both monkeys (F dot-dashed; I solid line) for (E) the adaptive exploration-exploitation value β*^RL^, (F) the error trace weight α ^ErrorTrace^, (G) the working memory softmax constant β^WM^ and (H) the strategy weight 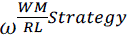. Stars in panel (C-H) denote standard significance levels.

We found that all three model mechanisms contributed to accounting for choices made by the subjects during task performance. This result is based on evaluating all combinations of model mechanisms (**Figure 5B**), which showed that the behavior across the stimulation and sham blocks of each monkey and in each area (ACC and striatum) is best accounted for by the full *AE-SF-WM model* combining all three learning mechanisms as evaluated with the Akaike Information Criterion (AIC) that corrects model performance for the number of free parameters used (**Figure 5B**; AIC’s for models rank-ordered from worst to best: (7) only adaptive exploration AE, (6) AE+WM, (5) only selective forgetting SF, (4) AE+SF, (3) only WM, (2) WM+SF, and for the best ranked model (1) SF+AE+WM for monkey 1 in ACC: 3138.9 / 3119.9 / 3024.5 / 3010.2 / 2998.7 / 2998.6 / 2967.5; for monkey 2 in ACC: 4819.6 / 4738.5 / 4596.3 / 4574.9 / 4525.6 / 4521.3 / 4491.6; for monkey 1 in striatum: 3492.5 / 3478.8 / 3496.5 / 3580.9 / 3489.9 / 3494.5 / 3744.0; for monkey 2 in striatum: 3331.7 / 3329.1 / 3332.8 / 3426.6 / 3329.5 / 3333.7 / 3573.7).

We next fitted the full *AE-SF-WM model* separately to the performance of stimulation and sham blocks to evaluate which model parameter values were affected by microstimulation. Microstimulation in ACC and the striatum affected similar model parameters, impairing (in ACC) and improving (in striatum) learning through changes of adaptive exploration parameters (α ^ErrorTrace^, ω^Exploration^), and RL parameters (learning rates for prediction errors), but less consistent or no changes in WM, and no changes of selective forgetting (**Figure 5C,D**). In particular, for the ACC, microstimulation changed parameters in each subject that implement adaptive exploration, altered RL learning rates, and increased WM/RL strategy weighting (**Figure 5C**, t-tests, Bonferroni corrected significance for monkeys F / I in ACC: adaptive exploration weight *ω^Exploration^*: 6.4 / 19.1, p<0.001 / p<0.001; RL learning rate for correct outcomes: t-stat: –15.5 / – 1.9, p<0.001 / ns; RL learning rate for incorrect outcomes: t-stat: 3.2 / 11.3, p=0.0199 / p<0.001; AE error-trace weight *α^ErorrTrace^*: t-stat: –3.9 / –5.3, p=0.002 / p<0.001; exploration-exploitation level β^∗*BL*^ t-stat: 4.6 / 20.7, p<0.001 / p<0.001; 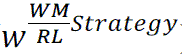: t-stat: –4.5 / –5.0, p=0.001 / p<0.001). The selective forgetting parameter value was unaffected by stimulation (**Figure 5C**; *ω****^selective For getting^*** for monkey F / I: t-stat: 0.9 / –0.5, ns / ns). The WM parameters for temporal decay, capacity and the exploration-exploitation level were not modulated or only moderately affected by stimulation in one of the subjects (**Figure 5C**; for monkey F / I: WM decay: t-stat: 2.2 / –3.5, ns / p=0.007; exploration-exploitation level β*^WM^* t-stat: –3.2 / –3.5; p=0.002 / p=0.008; WM capacity: t-stat: 1.8 / 1.6; ns / ns).

A similar set of model mechanisms was modulated by stimulation in the striatum. For each subject, striatal stimulation affected the adaptive exploration parameter values and RL learning rates for correct outcomes (**Figure 5D**; for monkeys F / I: adaptive exploration weight *ω^Exploration^*: t-stat: 19.3 / –1.01, p<0.001 / ns; RL learning correct outcomes: t-stat: 27.0 / 6.0, p<0.001 / p<0.001; RL learning rate after error outcomes: t-stat: 0.76 / –2.53, ns / ns; exploration-exploitation level β^∗*RL*^: t-stat: –29.7 / –6.95, p<0.001 / p<0.001). In contrast, WM parameters, selective forgetting and the strategy weight were either unaffected or inconsistently modulated across subjects (for monkeys F / I: β*^WM^* t-stat: –0.35 / 0.13, ns / ns; WM capacity: t-stat: 0.3 / –0.27, ns / ns; WM decay: t-stat: 5.05 / 0.14, p<0.001 / ns; *ω^SelectiveFor getting^*: t-stat: –0.33 / –13.18, ns / p<0.001; 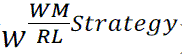: t-stat: –1.92 / 4.2, ns / p<0.001) (**Figure 5D**).

Both, the ACC and the striatum modulated adaptive exploration and the updating of feature values by scaling prediction errors with learning rates, but they did so in opposite ways. Stimulation in ACC increased exploration (lower β^∗^*^RL^*) and reduced weighting of recent errors to adapt exploration (*α^ErorrTrace^*) (**Figure 5E,F**), while in the striatum it enhanced exploitation (higher β^∗*RL*^), i.e. the use of expected values to make a choice, which was linked with a stronger weighting of recent prediction errors (*α^ErorrTrace^*) in subject F (**Figure 5E,F**), and to stronger weighting of exploitation over exploration in subject I (*ω^Exploration^*) (**Figure 5D**). These changes occurred while the model parameter that determined how strong the WM representations informed choices (β*^WM^*) remained unchanged in both subjects and brain areas (**Figure 5G**). While there were moderate increases of the strategy weight, 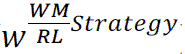, with microstimulation, the weight values of ∼0.2 reflected an overall low contribution of WM over RL representations to inform choices in both, stimulation and sham conditions (values of 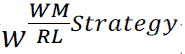∼ 0.5 would reflect equal weighting of WM and RL, whereas values<0.5 reflect less weight of the WM module), showing that stimulation did not change the dominant RL learning strategy (**Figure 5H**).

Taken together, microstimulation affected AE and RL model mechanisms in the ACC and the striatum (visualized in **Figure 6A,B**). To discern the consequences of changed model mechanisms on behavior we calculated the trial-by-trial changes of behavior (**Figure 6C,D**), and how the model variables of the *AE-SF-WM model* related to them. In experimental blocks with microstimulation (compared to sham) choice probabilities were reduced in ACC, but enhanced in the striatum (**Figure 6E,F**), prediction errors remained elevated in ACC, but were more effectively reduced in the striatum (**Figure 6G,H**), and subjects showed consistently more random exploration across trials with stimulation in ACC, but more effective exploitation with stimulation in the striatum (**Figure 6I,J**). A different pattern was found for the choice strategy, which temporarily increased for a few trials at the beginning of a block with stimulation in the ACC but stayed at a low (<0.5) level and thereafter remained largely unchanged in ACC and was unaffected by stimulation in striatum (**Figure 6K,L**).

**Figure 6.**
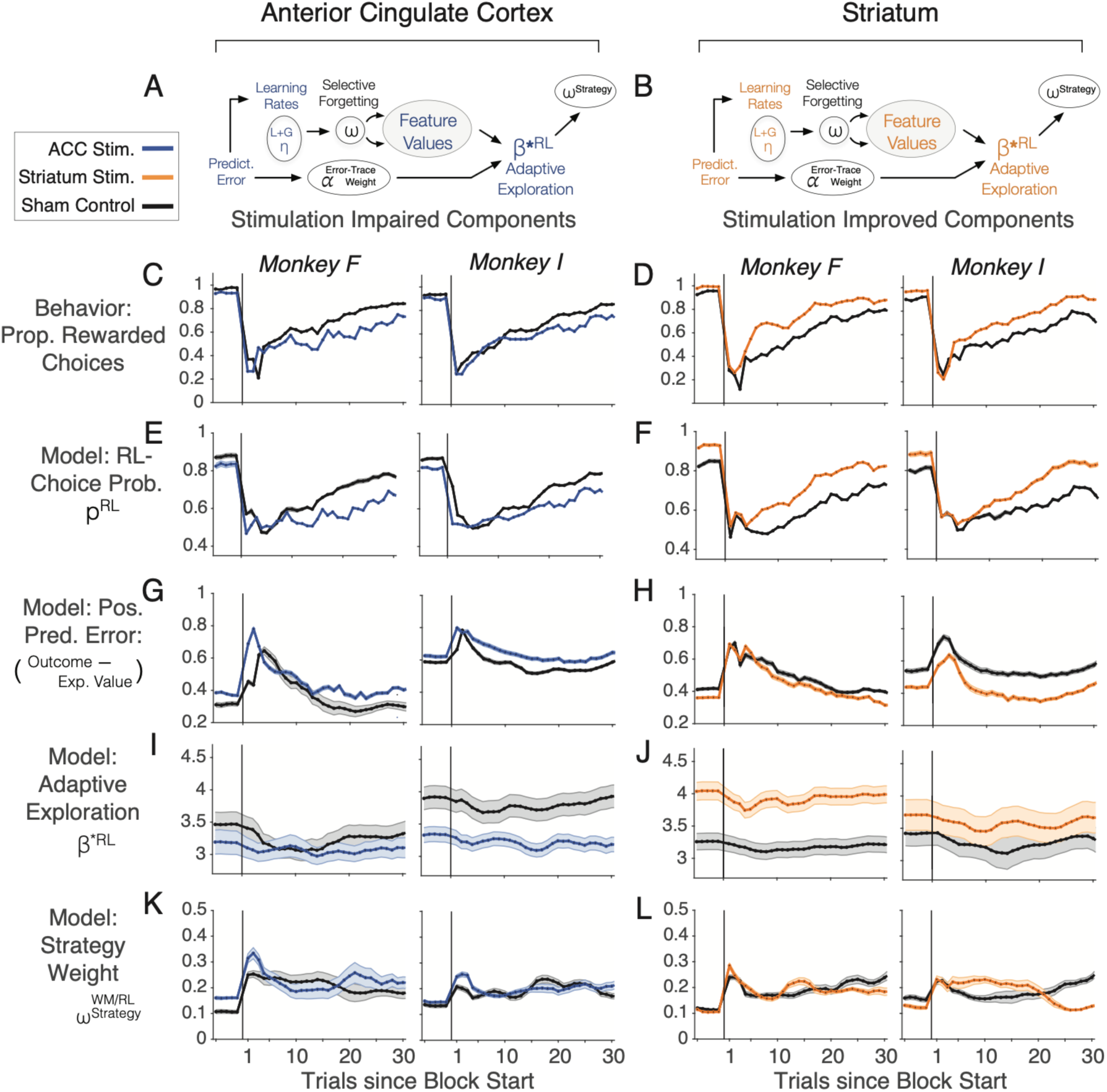
Stimulation has opposite effects in ACC and striatum on reinforcement learning and adaptive exploration. (**A,B**) Visual summary of the model mechanisms with coloring reflecting modulation with microstimulation in the ACC (*A*) and the striatum (*B*). See Figure 5A for the full model. (**C,D**) Trial-by-trial changes in behavioral accuracy for each monkey in ACC (*C*) and the striatum (*D*) when monkeys learned choices in the 3D condition in sham blocks (black lines) and stimulation blocks (colored lines). Analysis used the seventeen ACC sessions and seventeen striatum sessions in which stimulation affected behavior. (**E-L**) Same format as (*A,B*) showing estimated values from the AE-SF-WM model for the choice probability of the RL component (*E,F*), the positive prediction errors (*G,H*), the adaptive exploration-exploitation value β*^RL^(*I,J*), and the strategy weight 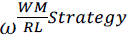 (*K,L*). Error shading is SD across fifty bootstrap model fits.

### ACC and striatum uses different neural codes for tracking errors and uncertainty

What are possible neurophysiological sources for the opposite effects of microstimulation on learning performance and model mechanisms in ACC (impaired learning, AE and RL), and the striatum (improved learning, AE and RL)? To address this question, we recorded neuronal activity with linear 64 channel silicon probes in subject F in ACC (737 recording sites) and in the striatum (835 recording sites) at those anatomical coordinates where stimulation had altered behavior (**Figure S7**). We fitted the behavioral performance with the *AE-SF-WM model* and evaluated whether activity in the ACC and the striatum differently encoded the learning variables of the model during the choice period (0.3-0.7 s following fixation onset) at which electrical stimulation was administered in the previous experiment. We found that neuronal activity in the ACC and the striatum showed different patterns of neural encoding of (*1*) choice probabilities, (*2*) object values, (*3*) trial outcomes and (*4*) error history (**Figure 7**). Firstly, there were more recording sites in the ACC that showed negative correlations of neural activity with choice outcomes during the choice, reflecting elevated activity that predicted that the forthcoming choice will be incorrect (unrewarded), while in the striatum there were more positively correlated sites with stronger firing during correct choices (**Figure 7A-C**; two-prop Z-test for neg. versus pos. correlations with correct/incorrect outcome during the 0.3-0.7 s in ACC: Two prop. Z-stat, –3.75, p = 0.00017; in striatum: Two prop. Z-stat, 3.1417, p = 0.00168; two-prop. Z-test for a difference in proportion of ACC vs. striatum for pos. correlations: Z-stat, –3.85, p = 0.00012; for neg. correl. Z-stat, 5.9501, p = 2.6798e-09). A similar pattern was evident for trial-by-trial correlations of firing with choice probability (**Figure 7D-F**) and with expected stimulus values (**Figure 7G-I**), both were being encoded by a larger neural population with negative than positive firing correlations in the ACC, and by a larger population with positive than negative correlations in the striatum (for choice probability: two-prop Z-test for neg. versus pos. correlations in ACC: Two prop. Z-stat, –4.7513, p = 2.0212e-06; in striatum: Two prop. Z-stat, 1.8402, p = 0.06574; two-prop. Z-test for a difference in proportion of ACC vs. striatum for pos. correlations: Z-stat, –1.2605, p = 0.20749; for neg. correl. Z-stat, 8.1773, p = 2.2204e-16; for the expected value of chosen objects: two-prop Z-test for neg. versus pos. correlations in ACC: Two prop. Z-stat, –1.9074, p = 0.05647; in striatum: Two prop. Z-stat, 4.3509, p = 1.3558e-05; two-prop. Z-test for a difference in proportion of ACC vs. striatum for pos. correlations: Z-stat, –3.4253, p = 0.00061; for neg. correl. Z-stat, 5.3876, p = 7.1387e-08) (**Figure 7F,I**).

**Figure 7.**
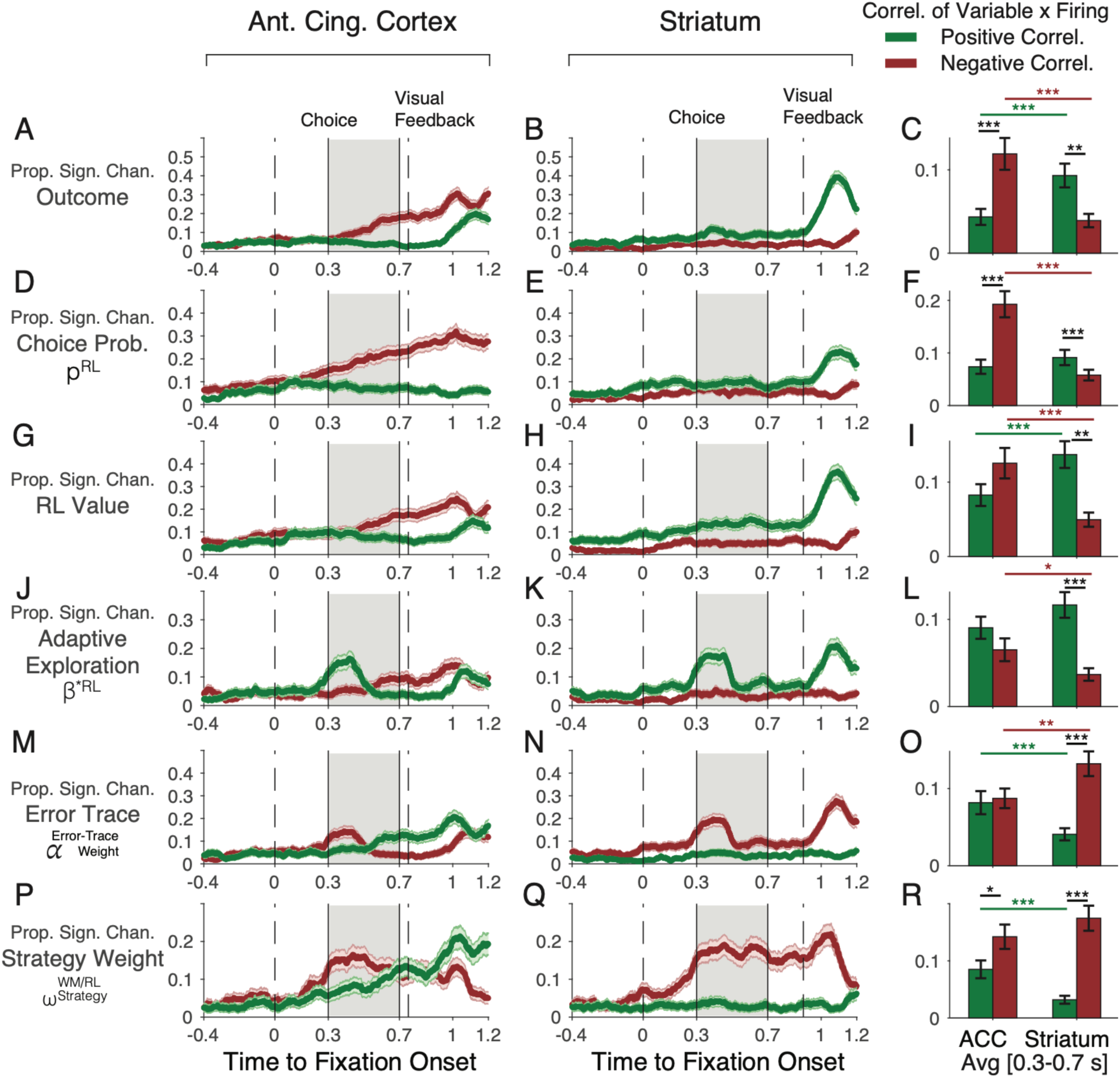
Encoding of behavioral and latent model variables with neuronal activity differs in the ACC and striatum. (**A,B**) The proportion of recording channels with multiunit activity in the ACC (*A*) and striatum (*B*) significantly correlating trial-by-trial with outcomes (0=unrewarded,1=rewarded) relative to the onset of fixating the chosen object. Green lines denote the proportion for channels with positive correlations, red lines denote proportions of channels with negative correlations. Grey shaded area is the time period when the subjects committed to choosing the fixated object (at 0.3 s) and when the choice is registered (0.7 s). This choice period was used for electrical stimulation. MUA activity was smoothed with a 0.2 gaussian window prior to the correlation with the variables. Correlations were performed every 25 ms. (**C**) Average proportion of channels with significant positive (green) and negative (red) correlations in ACC (*left*) and the striatum (*right*). Error bars are SDs, horizontal bars and stars denote proportional difference significance level. (**D-R**) Same format as *A-C* for neural activity correlations with choice probability of the chosen object (*D-F*), the expected values of the chosen object (*G-I*), the adaptive exploration-exploitation variable β*^RL^ (*J-L*), the error-trace weight variable (*M-O*), and trial-by-trial fluctuations with the WM/RL strategy weight (*P-R*).

For other variables, the coding in ACC and striatum was only partially different. The ACC and the striatum encoded the adaptive exploration parameter value in neuronal populations showing positive activity correlation, reflecting increased neural firing when there was more exploitation than exploration, i.e. when the adaptive exploration variable value was larger. The positive correlation was indistinguishable between areas (two-prop. Z-test for a difference in proportion of ACC vs. striatum for positive correlations: Z-stat, –1.6938, p = 0.0903) (**Figure 7J-L**). But in ACC, an additional population with negative firing correlations emerged during the choice period that was absent in the striatum (two-prop. Z-test for a difference in proportion of ACC vs. Striatum for neg. correl. Z-stat, 2.5747, p = 0.010032) (**Figure 7J-L**). These negatively correlated sites fired stronger when the choice was based more on exploring options than exploiting values, i.e. when the adaptive exploration variable value was lower. Similarly, a separate neuronal population in ACC was evident for the encoding of the error-trace that reflected how previous errors are accumulated to adjust values of the adaptive exploration variable. During the early choice period ACC and the striatum had a neuronal population with increased activity when errors accumulated, i.e. they were negatively correlated with the error-trace variable (two-prop Z-test for neg. versus pos. correlations with error-trace in ACC: Two prop. Z-stat, –0.27286, p = 0.78496; in striatum: Two prop. Z-stat, –4.7002, p = 2.5997e-06; two-prop. Z-test for a difference in proportion of ACC vs. striatum for neg. correl. Z-stat, –2.823, p = 0.0047575) (**Figure 7M-O**). But only the ACC had a separate neuronal population that increased firing with increasing error-trace values, evident in a positive firing correlation starting ∼0.5 s during the choice (two-prop. Z-test for a difference in proportion of ACC vs. striatum for pos. correlations: Z-stat, 3.416, p = 0.00063552) (**Figure 7M)**.

Similar to the error-trace correlations, the ACC and the striatum showed more often negative firing correlations with the strategy weight (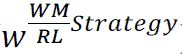), reflecting higher activity when the RL strategy dominated, i.e. when 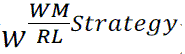 had a smaller value (two-prop Z-test for neg. versus pos. correlations with the WM/RL strategy weight in ACC: Two prop. Z-stat, –2.4396, p = 0.014702; in striatum: Two prop. Z-stat, –6.7735, p = 1.2571e-11; two-prop. Z-test for neg. correl. Z-stat, –1.7512, p = 0.079914) (**Figure 7P-R**). In addition to this negative correlation a neural population gradually emerged in ACC during the choice period with activity correlating positively with the strategy weight and which was absent in the striatum (two-prop. Z-test for a difference in proportion of ACC vs. striatum for pos. correlations: Z-stat, 4.5459, p = 5.4707e-0) (**Figure 7P,Q**).

## Discussion

We have shown that microstimulating the ACC and the striatum modulated the speed of learning the value of object features when there were multiple distracting features present (3D condition). Microstimulation in ACC and the striatum caused largely opposite behavioral effects. In the ACC stimulation impaired learning (**Figure 2C**), and this impairment (*1*) was more pronounced with higher motivational saliency when gains were higher (**Figure 2E,G**), (*2*) was more apparent following correct trials (**Figure 4A,C**; **S4K,M**), (*3*) was linked to a difficulty in sustaining correct responses (**Figure S6E**), and (*4*) gradually emerged with increasing stimulation trials (**Figure S6A**). In the striatum, stimulation improved learning (**Figure 2C**), and this improvement (*1*) did not increase with higher incentives (**Figure 2D-G**), (*2*) was more apparent following correct trials (**Figure 4A,C**; **S4O,Q**), and (*3*) was evident already following the first stimulation trials (**Figure S6C,D**). Beyond these overall effects, there was heterogeneity of the stimulation effects, but for each area, these overall effects were reliably evident in about a third of the stimulation sessions in ACC (monkey F / I: 29% / 42% of sessions) and in the striatum (monkey F / I: 33% / 38% of sessions) (**Figure 3**).

Taken together, these results document a dissociation of stimulation-induced changes of adaptive goal-directed behavior in the ACC versus the striatum. This dissociation became evident when the behavioral goal involved learning feature values in a task environment with multiple distracting features (3D condition). The impairment (in ACC) and improvement (in the striatum) of learning in this multidimensional environment was closely associated with a weakening (in ACC) and strengthening (in the striatum) of adaptive reinforcement learning mechanisms, while cognitive learning strategies such as selective forgetting and working memory were not consistently affected. The differences in behavioral stimulation effects were consistent with a more heterogeneous neuronal encoding of choice probabilities, values, error history, and adaptive exploration in the ACC when compared with the striatum (**Figure 7**). While both areas encoded these variables with firing rate increases (positive correlations of variables and firing), the ACC contained larger neuronal subpopulations than the striatum that showed negative firing correlations with choice probabilities, values, and exploration, while having separate neuronal populations that increased firing when errors accumulated (**Figure 7D,G**,**J,M**). This finding suggests that an activating electrical stimulation protocol (*see* below) will likely have increased neuronal signaling in the ACC linked to higher choice uncertainty, lower valuation of the chosen target stimulus, more exploratory choices and larger error traces. In contrast, activating the striatum will have more likely increased activity of neurons who would otherwise increase their firing when the choice was made with larger certainty (high choice probability, **Figure 7E**), to an object with higher value (higher value of the chosen stimulus, **Figure 7H**), and more exploitative choices (positive correlation with the adaptive exploration parameter, **Figure 7K**). These differential effects of stimulation are putative as they were collected at the same anatomical coordinates as the stimulation experiments, but not at precisely the same spot as they were not collected in the same experimental sessions. Despite this limitation, the consistency of differential stimulation induced behavioral effects, and the differential neuronal encoding provides a parsimonious framework for understanding our key results.

### Electrical microstimulation biases excitation of the local circuit output

An assumption for interpreting the main findings of our study is that the brief 0.3 s microstimulation pulse trains used in our experiment activated the neural circuit. Previous studies support this assumption, showing that biphasic ∼50 μA microstimulation pulse trains activate neurons within a ∼200-300 μm radius and increase net excitation across distances that can exceed 1500 μm, depending on the spread of excitatory horizontal projections ^52–55^. Microstimulation protocols with similar parameters (60-300 ms long, 20-100 μA pulses) applied in the frontal eye field and lateral intraparietal area bias saccades towards target stimuli overlapping with the peripheral response field of the stimulation site ^56–60^. Similarly ∼50μA microstimulation pulse trains have different effects in more mediodorsal frontal areas such as the supplementary eye field where microstimulation elicits eye movements at higher thresholds, can slow-down overt saccadic eye movements to target stimuli in difficult tasks, speed-up eye movements in simple tasks, and reduce the correct selection of sequential targets ^61–63^. These effects suggest that microstimulation in the supplementary eye field can enhance a net inhibitory effect of it on other saccade generating structures. These results suggest that stimulation in the non-eye movement related ACC (our stimulation sites were anterior to the eye movement fields in the mid-cingulate ^33^, **Figure 1D**) likely excites the local circuit, which is associated with an inhibitory effect on prefrontal projection targets ^64^ and a net excitatory effect on the striatum ^35^.

### ACC provides error monitoring and directed exploration over exploitation

Our stimulation effects provide evidence that the ACC is causally involved in the monitoring of errors and outcome uncertainty and in the directed exploration of visual objects to reduce outcome uncertainty during learning. Behavioral learning is governed by latent variables that are not explicitly observable but need to be estimated from the sequence of behavioral choices. We modeled various latent variables implementing learning mechanisms and cognitive strategies and found microstimulation affected similar mechanisms in ACC and striatum but in opposite ways (**Figure 5C-H**). ACC stimulation reduced weighting of recent errors (α ^ErrorTrace^). In the model lower weighting of error increases exploration (lower β^∗@A^ values). Consequently, the choices of both monkeys during ACC stimulation blocks reflected higher exploration levels (**Figure 6I**), more choice uncertainty (**Figure 6E**), and more outcome uncertainty as evident in higher prediction errors across the learning block in stimulation than in sham blocks (**Figure 6G**). The increases of uncertainty and exploration levels are consistent with the result that ACC stimulation induced a difficulty in sustaining corrected responses (**Figure S6E**), an effect that has also been found after extensive lesions of the ACC ^11^.

These results provide causal evidence that one overarching function of the ACC is to evaluate outcome uncertainty and to use this information to guide information seeking, i.e. to determine how to explore options to reduce uncertainty ^9,65,66^. In our study, these functions were affected in the 3D condition in which uncertainty is reflected in the number of features that could be associated with reward. In the presence of this larger feature space, subjects do not only need to know when to explore versus exploit, but also which specific feature should be credited for a choice outcome. Such a feature specific credit assignment is encoded in neurons in the ACC during feature learning ^29^, and these neuronal representations of feature-specific prediction errors provide critical information to guide exploration towards those features that result in positive prediction errors and to avoid choosing features whose choice results in negative prediction errors.

Taken together, our modeling results suggests that the ACC contributes to an adaptive learning process by tracking the history of recent prediction errors to adjust the degree of exploration over exploitation that subjects apply in subsequent trials. This adjustment of exploration may be either un-directed, i.e. modifying random exploration, or directional, i.e. modifying which features to explore in future trials. Our results are more consistent with the ACC supporting directed exploration, because stimulation did not induce random exploration in the 2D or 1D condition but specifically altered exploration in the 3D condition when high feature uncertainty calls upon exploratory guidance. In this 3D environment learning was impaired most prominently following gains compared to losses – irrespective of whether stimulation was administered on correct or error trials – suggesting that assigning positive credit to specific features for exploratory guidance was disrupted more than using loss to avoid features that were unrewarded. These conclusions resonate with a hypothesis that posits the ACC acts together with the ventrolateral prefrontal cortex to guide information seeking by detecting the sources of uncertainty and directing exploratory sampling towards objects to reduce the uncertainty ^65^. This hypothesis also clarifies that the role of the ACC for monitoring sources of uncertainty requires interactions with the ventrolateral prefrontal cortex to update those representations that guide information seeking, attention, and choices, consistent with lesion studies ^67,68^, and with recording studies showing neuronal tuning to attended object features ^69,70^ and attention-dependent spiking interactions between the ACC and the ventrolateral prefrontal cortex ^35,71^.

### Causal facilitation of adaptive reinforcement learning with striatum stimulation

In the striatum, the microstimulation effects on behavior and on the learning mechanisms of the learning model were opposite in sign compared to the ACC. Striatum stimulation enhanced exploiting expected values over exploration (higher β^∗RL^), and improved the adaptive reinforcement learning mechanisms either by enhanced weighting of recent errors (α ^ErrorTrace^, in one subject) (**Figure 5E,F**), or by an enhanced exploration weighting (ω*^Exploration^*, in the other subject) (**Figure 5D**). These changes were reflected in reduced prediction errors (**Figure 6H**), enhanced certainty (choice probability) with which choices were made (**Figure 6F**), and more exploitation of value expectations (**Figure 6J**). These findings provide evidence that the striatum causally supports adaptive reinforcement learning. This conclusion is consistent with suggestions that the striatum is situated in fronto-striatal networks to adjust learning processes about goals versus values ^10,72^, and about predictive versus selective value associations ^1^. However, causal evidence for a role in adjusting reinforcement learning during goal-directed behavior has remained sparse. Our findings indicate that striatal microstimulation enhances the weighting of prediction errors, which will improve value updating and reduced uncertainty about the most valuable feature in future trials. Support for this scenario comes from multiple sources. Previous studies have found that striatal neurons that encode feature-specific prediction errors begin to encode the value of reward predicting stimuli when learning completes ^29^, and that subpopulations of striatal neurons fire stronger the less uncertain the reward-value of attended stimuli ^28,29^. These findings were confirmed in our neural recordings that revealed larger populations of neurons encoding choice probability and the value of chosen stimuli with firing rate increases (i.e. via positive correlations of firing with variables) than with firing decreases (**Figure 7E,H**). In addition, a prior stimulation study has documented that stimulation of the caudate nucleus when subjects process a single stimulus on the screen can enhance the value of that stimulus, effectively enhancing the likelihood it will be chosen in future choices ^31^. Our results extend these findings by suggesting that stimulation does not enhance the value of all features of a processed object, but that stimulation selectively enhances the reward-relevant target feature particularly when there were two distracting features of the multidimensional objects that need to be filtered during learning. The specificity of the stimulation effect to improve learning in the 3D condition is consistent with the overarching function of the striatum as implementing an adaptive filtering of cortical inputs. This filtering function of the striatum has largely been inferred for the motor domain, but recent studies have started documenting that similar adaptive filtering and attentional selection is evident also for non-motor tasks that involve the learning and choices of visual features and objects ^28,35,42,73,74^.

### ACC mediates cognitive control at high motivational saliency

We observed the most apparent learning deficits with ACC stimulation when two conditions were met: when the motivational salience of a choice was high (high gains or losses) and when the credit assignment about feature values was difficult (in the 3D condition). This pattern of result is consistent with a study causing a more prolonged disruption of the ACC ^17^ and shows that the ACC fundus multiplexes motivational and cognitive functions, which is consistent with two major accounts of ACC function: Consistent with a motivational effort-control account ^75^, the ACC causally supported learning when the motivational saliency of choices is high by either anticipating a high motivational benefit (gaining 3 vs 2 token) or (in one subject) a high potential punishment (loss of 3 vs 1 token). Consistent with a cognitive account, the ACC causally guided searching for choice options when outcome uncertainty was high (3D versus 1D or 2D) ^19,76^. Importantly, the multiplexing of motivation and cognitive variables is consistent with the ACC’s rich direct anatomical connectivity to subcortical sites that directly affect motivation, including dopaminergic neurons, and that mediate attentiveness and cognitive control, including cholinergic and norepinephrinergic nuclei ^77,78^. This specific anatomical connectivity has been used as a key argument for suggesting that the ACC adaptively boost cognitive control when task demands are high ^79^. Our results provide direct causal support for this ‘boosting’ function of the ACC. Importantly, our modeling results suggest that the proposed theoretical function of the ACC to ‘boost cognitive control’ ^80^ may be realized by modulating exploration when a difficult task requires motivational support ^17^.

Taken together, the present study reveals a causal influence of the ACC and the striatum to support adaptive reinforcement learning. The behavioral stimulation effects were traced back to the ACC’s and striatum’s role in the ongoing estimation of the reinforcement learning values and their uncertainty given the history of recent experiences and to the feature-specific credit assignment that is needed to optimize attention and decision making in a changing environment. These results and conclusions may prove important for developing brain computer interfaces designed to enhance cognitive control and support cognitive flexibility.

## Methods

### Ethics Statement

All animal and experimental procedures were in accordance with the National Institutes of Health Guide for the Care and Use of Laboratory Animals, the Society for Neuroscience Guidelines and Policies, and approved by the Vanderbilt University Institutional Animal Care and Use Committee (M1700198-01).

### Experimental Procedures

Two adult male macaque monkeys performed the experiments (monkey F, 13 years of age, 11.3 kg, and monkey I, 11 years of age, 11.0 kg). Behavior and electrical stimulation triggers were controlled using the Multi-Task Unified Suite for Experiments (M-USE) ^81^. The animals were trained on the feature-reward rule learning task in the cage-based touchscreen Kiosk Testing Station ^82^ and stimulation experiments proceeded in sound attenuating experimental booths with subject’s head position fixed, facing a 21’’ LCD screen at a distance of 63 cm from their eyes to the screen center. For each stimulation experiment, a tungsten stimulation electrode (FHC Inc., Bowdoin, ME) was lowered through a guide tube at a pre-determined location in a custom recording/stimulation chamber implanted over the left hemisphere. The stimulation electrodes were lowered into the ACC or the striatum and stopped once sustained spiking activity was noticeable and constant.

### Task paradigm

The feature-reward rule learning task required learning through trial-and-error which feature of multidimensional objects is associated with reward. Objects were 3D rendered Quaddles ^83^ that could vary features in 1,2, or 3 feature dimensions within a block (**Figure 1A-C**). The feature dimensions were body shape, arm type, color, and body pattern with 9-11 feature per dimensions (e.g. oblong, pyramidal, spherical, etc. for body shapes). In each block, only one feature value was associated with token reward. The feature that was rewarded, i.e. the feature-reward rule, stayed constant for 30-50 trials, then switched randomly to another feature in a new block. Monkeys initiated a trial by fixating a black circle for 0.5 s, which triggered after a 0.3 s delay the presentation of three objects randomly positioned at three of the four corners of a virtual square-grid spanning 10.5 cm with 5° eccentricity relative to the screen center. Objects extended over ∼3cm on the screen corresponding to ∼2.5° degrees of visual angle. The animals had up to 5 s to choose one object by maintaining gaze at an object for >0.7 s Choosing the correct object was followed by a yellow halo around the stimulus as visual feedback, an auditory tone, and either 2 or 3 tokens (green circles) added to the token bar (*G2* and *G3* conditions). Choosing an object without the rewarded target feature was followed by a blue halo around the selected objects, a lower-pitched auditory feedback, and for the loss conditions, the presentation of one or three grey ‘loss’ token that traveled to the token bar where one or three already obtained tokens were removed. In different blocks either 1 or 3 tokens were lost after incorrect choices (*L1* and *L3* conditions). The timing of the feedback was identical for gain and loss conditions. In each session, monkeys were presented with up to 36 separate learning blocks, each with a unique feature-reward rule.

Across all 48 experimental sessions, monkeys F/I completed on average 33.5 / 36 blocks during ACC stimulation sessions and 32.0 / 35.8 blocks during striatum stimulation sessions. For each experimental session, a unique set of objects was defined by randomly selecting three dimensions and three features per dimension (e.g., 3 body shapes: oblong, pyramidal and ellipsoid; 3 arm types: upward pointy, straight blunt, downward flared; 3 patterns: checkerboard, horizontal striped, vertical sinusoidal) ^83^. Based on this feature set three different task conditions were defined: One task condition contained objects that differed in features of only one dimension, while features of the other dimensions were identical; for example, the object body shapes were oblong, pyramidal, and ellipsoid, but all objects had blunt straight arms and uniform grey color (‘1D’ condition). A second task condition defined objects that varied features in two dimensions (‘2D’ condition), and a third task condition defined objects that varied in features of three dimensions (‘3D’ condition). Learning which feature is rewarded is systematically more demanding with objects varying in more feature dimensions ^17^.

### Experimental design

Each session was composed of 36 feature-reward rule blocks that pseudo-varied four motivational conditions and three cognitive demand conditions. Cognitive demand varied the number of object features that varied from trial to trial with features varying in one, two, or three dimensions (conditions 1D, 2D, or 3D). The motivational conditions varied the amount of tokens gained for correct choices to be 2 or 3 (G2 and G3 conditions) and the amount of tokens subtracted (lost) from incorrect choices were either –1 or –3 (L1 and L3 conditions) as introduced in prior studies ^84^. Subjects had to earn 8 tokens to completely fill a token bar on top of the screen. Each block began with an asset of 3 tokens. This starting asset ensured that subjects lost tokens already after the first incorrect trial in a block and thereby recognized the block’s loss condition. Across the 36 blocks all combinations of motivational conditions (+2/-1, +3/-1, +2/-3, +3/-3) and cognitive conditions (1D, 2D, 3D) were pseudo randomly assigned to blocks in equal number.

Each session contained 15 blocks with electrical stimulation, 15 sham stimulation blocks, and 6 initial blocks without stimulation. A session was designated either a *StimReinf+* or a *StimReinf-* session in which *Sham* and one of the stimulation conditions (*StimReinf+* or *StimReinf-*) alternated starting after the sixth block. A session ended either when all 36 blocks were completed, or when a monkey stopped initiating trials after 90 minutes.

### Monitoring and analysis of gaze

Gaze was monitored with the binocular infrared eye-tracker at 600 Hz sampling rate (Spectrum, Tobii). Gaze was calibrated before the task began with a 9-point eye-tracker calibration routine. Object fixations were reconstructed using a robust gaze classification algorithm described in ^85^. The duration of object fixations was analyzed for objects fixated prior to the last fixation onto the object that the subjects choose by maintaining fixation for >0.3 s onto that object.

### Electrical microstimulation

Electrical stimulation was delivered with the Intan Stimulation Controller (RHS 2000, Intan Technologies, LLC), starting 300 ms after monkeys continuously fixated an object and lasted for 300 ms. Choice fixations lasted >700 ms ensuring that the electrical stimulation was administered only while the chosen object was fixated. Monkey’s fixation durations had a bimodal distribution with the fixations indicating the sampling of information prior to making a choice lasting 150-200 ms on average in the 1D, 2D, and 3D conditions, and the last fixation that monkeys used to indicate a choice lasting on average 800 ms (see Suppl. Figure S7 in ^17^). The 300 ms lasting 200 Hz electrical stimulation pulse train consisted of 60 biphasic 50 μA pulses, with a 0.2 ms cathodal current injection followed immediately by a 0.2 ms anodal current, and a 5 ms interval between onsets of these 0.4 ms lasting pulses (*see* inset in **Figure 1A**). Prior to starting microstimulation in an experimental session, we measured the impedance using the built-in Intan stimulation controller’s *electrode impedance measurement*, resulting in an average impedance of 94.45 and 85.06 *k*Ω for stimulation sessions of Monkeys F and I. These values are as expected given the 50kΩ – 100kΩ impedance of the stimulation electrode (tungsten microelectrode UEWLFDSEAN1E, 100 mm length; FHC Inc., Bowdoin ME).

### Stimulation-triggered multi-unit activity and its relation to learning performance

Prior to starting stimulation within an experimental session, we ensured through visual and auditory inspection of neural activity that the stimulation electrode was in the grey matter at sites in which neurons showed spiking activity. Offline, we quantified how strong multi-unit activity at the stimulation electrode was modulated by electrical stimulation and tested whether the modulation of stimulation-triggered multi-unit activation (MUA) was predictive of behavioral effects of stimulation. To this end, we extracted continuous multi-unit activity using a 0.5-3 kHz bandpass filter, followed by rectification, and 200 Hz low-pass filtering of the recorded signal before and after the 0.3 s stimulation period. We removed outlier trials reflective of artifacts with the inter-quartile method and for each session normalized the data to a –0.3-0 s pre-stimulation baseline period. To quantify the stimulation triggered effect, we extracted the maximum response within a 0.65-1.3 s post-stimulation window in the stimulation and sham conditions, and calculated a *MUA Index* as the normalized difference (*Stim* – *Sham*)/(*Stim* + *Sham*) of that maximum response. The *MUA Index* ranges from –1 to +1 with positive values reflecting stronger MUA activity in the stimulation condition compared to the sham condition. For the same sessions we also quantified the learning speed (trials-to-criterion) and calculated a *Performance Index* (*Stim* – *Sham*)/(*Stim* + *Sham*). We statistically analyzed the relation of the MUA Index and the Performance Index for those experimental conditions in which stimulation affected learning using Pearson correlations.

### Reconstruction of anatomical stimulation locations

We reconstructed the recording and stimulation sites in the ACC and striatum using the software 3D slicer (http://www.slicer.org, version 4.11) (**Figure 1D,E**). For each monkey, we first co-registered MRI’s taken when the recording chamber was implanted, with CT scans that were taken after the chamber was implanted. Then, we segmented the recording chamber and reformatted the image space to reference the chamber space. We recorded the x,y,z (z being the depth of the electrode) coordinate the stimulation electrode tips relative to the chamber to reproduce the penetration depth for each stimulation electrode. The reconstructed electrode tip site was labeled on the MRI images, where monkey F’s stimulation sites were overlayed onto monkey I’s image to have a unified image of stimulation sites.

### Electrophysiological recording experiment

Following the stimulation experiment, we performed electrophysiological recordings in thirteen experimental sessions in the ACC and the striatum at those anatomical coordinates used during the stimulation experiment. Recordings were performed in monkey F performing the same experimental task as during the stimulation experiment. In these recording-only sessions neural activity was recorded with 64-channel linear silicon probes (Neuronexus, 50 μm interelectrode distance) using Open-Ephys (open-ephys.org) and the Intan Recording System (RHD2000, Intan Technologies, LLC). Neuronal wideband recordings were notch filtered (60, 120, and 180 Hz) with a filter bandwidth of 2 Hz to remove power noise and its harmonics and re-referenced using median referencing across channels. Continuous multi-unit activity (MUA) was quantified by 750-5000 Hz bandpass filtering, rectifying, 300 Hz low pass filtering, and down-sampling to 1000 Hz. Individual trials were extracted from trial initiation until the token and reward feedback was given with additional 0.5 s padding. Trials were aligned to the time of the decision, and MUA smoothed using a gaussian window of 0.2 s. We then used the mean activity during the baseline period 0.5 s at the beginning of the trial and prior to the onset of the stimuli to detrend the data channel-wise. We then determined whether a recording channel showed task modulated neuronal activity with a regression model that predicted MUA activity every 25 ms using as factors trial outcome (correct vs error), reward outcomes (gain and loss of tokens), learning status (before vs. after trial-to-criterion was reached), stimulus dimensionality (1-,2-, and 3 dimensional objects), and previous trial outcome (correct vs error). We removed channels that showed no task modulation for any of these task variables in at least three consecutive time windows which affected on average 0.5 channels per probe.

For the neuronal analysis, we aligned the MUA activity to the onset of the fixation of the object the subject chose on that trial by continuing fixating it for 0.7 s and smoothed it with a 200ms gaussian kernel (**Figure 7**). We then calculated every 25 ms the correlation of MUA activity across trials with the outcomes (error vs. correct) and with trial-by-trial changes of the model-estimated choice-probability, prediction error, expected object value of the chosen object, adaptive exploration value, error-trace, and the strategy weight (see Figure 7 and Methods). For each variable we quantified the proportion of MUA channels that shows positive versus negative correlation values at a p<0.05 significance level. We compared the average proportion of positive versus negatively correlations in the 0.3 – 0.7 s time window for each variable within each area and between areas (ACC versus Striatum) with a 2-sample Z test for proportions (**Figure 7C,F**,**I,L,O,R**). The time window overlaps with the 0.3-0.6 s time window that was used for electrical microstimulation.

### Behavioral analysis of learning and plateau performance

We estimated the efficiency of learning by calculating the trials required to reach criterion performance as the first trial within a block at which animals reached 70% accuracy over the following 12 trials. To compare the trials-to-criterion between conditions we used two-way Student’s T tests with FDR corrected (alpha = 0.05) p-values. If monkeys did not reach this learning criterion during a block, we calculated the trial at which the learning criterion would have been reached by using a linear regression of the last 12 trials in the block to find the predicted trials needed to reach criterion. We counted these blocks as being unlearned to determine that the proportion of learned blocks did not differentiate the experimental conditions (**Figure S1A**). For the condition with *StimReinf-* stimulation, we excluded blocks with less than 2 active stimulation trials (i.e. with less than 2 errors). We applied two-way Student’s T tests with FDR corrected (alpha = 0.05) p-values to compare plateau accuracy of the animals. Plateau performance was indexed as the average accuracy across trials once the learning criterion was reached (**Figure S1B**).

### Analysis of Variance (ANOVA) on experimental conditions

N-way ANOVAs were performed to determine if there were significant stimulation effects on learning (measured by trials-to-criterion for the block) between four experimental factors: dimensionality (*Dim*) with three levels (1D, 2D, and 3D); *Gain* context with 2 levels (Gain 2, Gain 3); *Loss* context with 2 levels (Loss 1, Loss 3), and stimulation (*Stim*) contexts 3 levels (Sham, *StimReinf+*, *StimReinf-*). We reported the significant (p < 0.05) interaction effects as either 2-way (stimulation condition x dimensionality) or 3-way (stimulation condition x dimensionality x gain/loss) effects.

### Identifying sessions with individually significant behavioral stimulation effects

We identified sessions in which stimulation impaired or improved learning relative to sham using Fisher’s Exact test comparing stimulation versus sham blocks in the 3D condition (**Figure 3**). We computed an index to facilitate comparison of the trials-to-criterion (learning speed) across sessions as *Index = (Stim – Sham)/(Stim-Sham)* with values ranging from +1 to –1, with positive values indexing slower learning in the stimulation than sham blocks and negative values indexing faster learning in the stimulation than sham blocks.

### Linear mixed effect models for block-level analysis of behavioral metrics

We used linear mixed effects models (LME) to evaluate the influence of experimental parameters on several behavioral metrics, such as 1) trials-to-criterion, labeled ‘LP’ for learning point; choice reaction times 2) during learning (prior to reaching the learning point), and 3) after learning; the number of fixations prior to choosing an object 4) during learning and 5) after learning; and 6) the plateau accuracy when learned completed. We tested stimulation effects on behavior with four experimental factors: feature dimension (*Dim*) with three levels (1D, 2D, and 3D); *Gain* context with 2 levels (Gain 2, Gain 3); *Loss* context with 2 levels (Loss 1, Loss 3), and stimulation (*Stim*) contexts with 2 or 3 levels (Sham, *StimReinf+*, *StimReinf-*). We used two other factors as random effects: the feature dimension of the target feature (*Feat*) with 4 levels (color, pattern, arm, and shape) and subject (*Monkey*) factor with 2 levels (F, I). The LME is formalized as *Metric* = *Dim* + *Gain* + *Loss* + *Stim* + (1|*Feat*) + (1|*Monkey*) + *b* + ε. We extended this LME to test for interactions of *Stim2* with *Dim*, *Gain*, and *Loss: Metric* = *Dim* + *Gain* + *Loss* + *Dim* × *Stim* + *Gain* × *Stim + Loss* × *Stim* + (1|*Feat*) + (1|*Monkey*) + *b* + ε.

### Analysis of choice reaction times

Stimulation might have an influence on the reaction time from display onset to the time when an object is fixated. We analyzed choice reaction times separately for correct trials in which the subjects made only one saccade and chose the first object they fixated and trials in which they fixated a second and third object (**Figure S2A,B**). The distribution of choice reaction times was determined separately for the sham, *StimReinf+*, and *StimReinf-* conditions from the first block in which stimulation began (the sixth block onwards). Choice reaction time distributions for trials with one, two, and three object fixations were compared with a two-way Student’s t-test with FDR corrected (alpha = 0.05) p-values.

### Analysis of fixational sampling of objects

To test whether stimulation altered fixational sampling of objects we analyzed the number of exploratory fixations of objects prior to the last choice-fixation during learning (prior to reaching learning criterion) and when learning completed for the sham and stimulation conditions, comparing them using permutation tests with FDR corrected (alpha = 0.05) p-values (**Figure S1E,F**). In addition, we compared the duration of stable fixations on objects prior to the last fixation of an object that indicated it was chosen (**Figure S2A-B**). Fixational sampling duration distributions for trials with one, two, and three or more object fixations were compared with a two-way Student’s t-test with FDR corrected (alpha = 0.05) p-values. The distribution of fixational sampling duration was done separately for the *sham*, *StimReinf+*, and *StimReinf-* conditions from the first block in which stimulation began (the 7^th^ block onwards).

### Temporal stability of stimulation effects on learning across sessions and within sessions

To discern whether the effect of stimulation on learning was stable over time within a session, we fitted a regression to the average trials-to-criterion for sham and stimulation blocks per session and compared the slope and intercept across conditions for each monkey, separately for the 1D, 2D, and 3D conditions (**Figure S2C,D,G,H**). To test for temporal stability of the stimulation effects within a session, we subdivided the number of stimulation blocks into three equally sized sets of ten blocks (blocks 7-16; 17-26; 27-36) and calculated the average trials-to-criterion criteria separately for the stimulation conditions and for the 1D, 2D, and 3D conditions (**Figure S2E,F,I,J**). We used the two-way Student’s T test with FDR corrected p-values (alpha = 0.05) to test for significance between the time epochs within a session.

### Analysis of the behavioral adjustment following erroneous and correct choices

We quantified the accuracy (proportion correct choices) for those trials that were preceded by different types of behaviors separately for the *Sham*, *StimReinf+*, and *StimReinf-* conditions. These analyses were limited to trials during learning, i.e. prior to completion of learning and performed with data combined from both monkeys for all sessions (**Figure 4**) and limited to the effective sessions with individually significant behavioral stimulation effects (**Figure S4K-R**). We separately calculated accuracy in trials relative to an erroneous or a correct choice. The main analysis was limited to the trials prior to completion of learning, but results were qualitatively similar when all trials were considered.

To compare stimulation effects on previous trial dependent performance, we used permutation tests. For the post-error and post-correct performance, we randomly shuffle, without replacement, the stimulation condition labels (sham, *StimReinf+*, *StimReinf-*) of the trial outcome for each n^th^ trial (n=1-5 trails following the outcome), for 10000 iterations to establish a randomized sampling distribution of the mean difference between groups. We then calculated the probability of a finding a value greater than the observed mean difference per stimulation condition following an outcome; the null hypothesis being that accuracy following a given trial outcome is not different across stimulation conditions. The p-value calculated is the probability that the difference between the groups mean being greater than the observed mean for each n^th^ trial. P-values were FDR corrected (alpha = 0.05) across post outcome trials.

### Accuracy relative to the number of stimulation trials

We tested how accuracy changed with increasing number of stimulation trials by calculating the average performance across the subsequent 3 trials following the 1^st^, 2^nd^, 3^rd^, … n^th^ stimulation trial within stimulation blocks and sham-stimulation trials in sham blocks (**Figure S6A-D**). We compared conditions using permutation tests for each subject and the 1D, 2D, and 3D experimental conditions.

### Error-Correct (EC_n_) analysis of trial effects on accuracy

We tested if stimulation affected the ability to sustain correct or incorrect choices over trials (**Figure S6E-H**). We calculated the accuracy in the trial following correct (C) and incorrect (E) sequences occurring in the block, where ‘CEE’ would correspond to a ‘Correct, Incorrect, Incorrect’ sequence the accuracy of the following trial. We calculated this across blocks and separately for the 1D, 2D, 3D conditions separately for the sham versus *StimReinf+* and sham versus *StimReinf-* comparison. The choice outcome sequences used for the Sustained Correct analysis were: E, EC, ECC, ECCC, ECCC; and for the Sustained Error analysis: C, CE, CEE, CEEE, CEEEE. P-values were FDR corrected (alpha = 0.05) for permutation testing.

### Logistic regression model to compare trial-wise differences between stimulation and sham

We used a generalized linear model (GLM) with a binomial distribution and logit link function to analyze binary choice outcomes across trials in sham and stimulation conditions. The model tested how trial number (*Trial*) following gains or losses, and stimulation condition (*Stim*) influenced response probabilities (correct = 1, incorrect =0), while also considering their interaction. Thus, the GLM was formalized as: *logit*(P(*Y*)) = β0 + β1(*Trial*) + β2(*Stim*) + β3(*Trial* × *Stim*) + ε, where *P(Y)* is the probability of a response (1 or 0), and ɛ represents residual error. To assess the significance of stimulation effects, we examined the estimated coefficients (β1, β2, β3) and their corresponding p-values (Wald tests). A significant interaction (β3) would indicate that the influence of trial number on response probability differs between sham and stimulation conditions (interaction). This approach allows us to statistically test whether stimulation alters behavioral response patterns over time. For analyses involving trials following gain or loss, only the five subsequent trials were included in the model.

### Behavioral modeling combining reinforcement learning, working memory and adaptive exploration

We modeled the choices of the monkeys with a recently proposed reinforcement learning model ^23^ that combines the classical mechanism of reinforcement learning (RL) of values using reward prediction errors with (1) an adaptive exploration (AE) mechanism that adjusts the degree of exploration relative to exploitation depending on the recent history of negative prediction errors (when a reward was predicted but not obtained), (2) a selective forgetting (SF) mechanism that decays the values of features from non-chosen (non-attended) objects, and (3) a working memory (WM) mechanism that updates objects in a working memory after a rewarded choice. The full model combining all three mechanisms is labeled the *AE-SF-WM model*, depicted in **Figure 5A**. It generates on every trial two hypothetical choices (choice probabilities) based on the RL and the WM component and arrives at the final choice by weighting the expected feature values (RL) and object values (WM) using a weighting parameter that reflects the relative success of each mechanism of predicting rewarded choices in the past trials (**Figure 5A**). We refer to this weighting parameter as the strategy weight as its value indicates whether the RL component or the WM component is contributing more to a choice. There are several static model mechanisms as well as two adaptive variables of the adaptive RL-WM model that vary trial-by-trial depending on the performance of the subjects. These model mechanisms have been systematically explored and validated to be important for adaptive behavior with multidimensional stimuli elsewhere ^23^. In brief, the reinforcement learning (RL) model updates expected feature values (V) using reward prediction errors (RPEs) scaled by learning rates. The RL component is a traditional Rescorla Wagner Q-learning model that estimates for each trial the expected Q value of objects. For each of the three objects presented on a given trial, the object value is calculated as the sum of the values of its features (V^F^), which are updated on every trial according to the reward prediction error (RPE) equal to (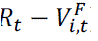), where R is the rewarded (1) or nonrewarded (0) outcome of the trials multiplied by a learning rate η*_t_* resulting in the equation 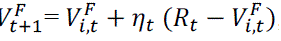, here *i* indexes the feature and *t* the trial. We evaluated models with two separate learning rates for error outcomes (η_t_ = η*_LOSS_*), and for correct outcomes (η*_t_* = η*_Gain_*), similar to Ref. ^86^. The RL component was augmented with a selective forgetting (SF) mechanism by decaying the values of non-chosen (nc) and not-presented (np) features by ωnc,t = ωnp,t = ω^selectiveForgetting^. This mechanism prioritizes learning from chosen object features, which are also those that were attended when making a choice, which makes the SF an attentional filtering mechanism that has been shown to be important in previous studies ^23,29,50^. On any given trial, each object has a probability of being chosen from the RL component of the model according to a softmax function applied to the object values. Specifically, the choice probability is calculated as 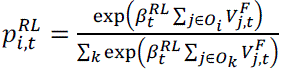, which uses a β^RL^ parameter (the so called *inverse temperature* that we interpret as exploration/exploitation weight and can vary from trial-to-trial in our model setting, see below) to weigh the sum of the values of all features (V^F^) of the presented set of objects (O_i_). At low β^RL^ the choice is exploratory and less strictly determined by the value of objects, while at high values of β^RL^ the choices follow the highest valued object and thus are exploitative.

*Fast one-shot working memory.* Working memory of object values is implemented using static parameters for the capacity and the decay of working memory content. An object will be represented in WM when it is chosen and rewarded, which is represented as an object’s WM value (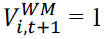) ^3,21^. WM values *V^WM^* decay slowly by *decay^WM^* which allows its maintenance over a few trials as long as the same object is not chosen without being rewarded. When an object is chosen and not-rewarded, *V^WM^* is reset to the default value 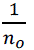, here n_o_ is the number of presented objects. This rapid updating of the WM after rewarded and unrewarded choices of an object reflects a ‘one-shot’ learning mechanism that does not use reward prediction errors (**Figure 5A**). WM content contributes to the choice according to a β*^WM^* parameter determining the choice probability *p^WM^* of object *i* on trial *t* in WM through a softmax selection process 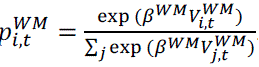.

*Adaptive changes of exploration-exploitation balance.* We use an adaptive exploration (AE) mechanism that adjusts the balance of exploration and exploitation. In the RL model with the AE mechanism, a choice is predicted by a β*^RL^ exploration parameter that is not fixed but varies over trials with the history of recent negative prediction errors. When subjects make successive errors (i.e. reward prediction errors (*δ_t_*) are negative) during learning a new feature-reward association the β*^RL^ value is reduced to increase exploratory choices among objects ^23,25^. In contrast, the β*^RL^ parameter is initially increased after correct trials (more exploitative) after which reward prediction errors start decreasing, hence the increase in β*^RL^ attenuates. The above behavior of the time-varying β*^RL^ is captured by weighting after each trial reward prediction errors with a magnitude that depends on its sign in terms of an auxiliary variable 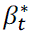. Specifically, 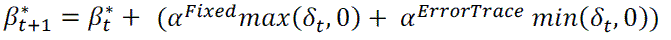, where α*^ErrorTrace^* is a free parameter to scale the error-trace (the α*^Fixed^* parameter is fixed, as only its relative size matters) and *δ_t_* is the reward prediction error computed across all object features (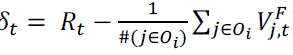), here i is the index of the chosen object and t the trial. Both α*^ErrorTrace^* and α*^Fixed^* are negative, which means a positive RPE decreases 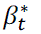 and a negative RPE increases it. This prediction error trace is translated into an actual β*^RL^ value by transforming it with a (negative) exploration factor ω^Exploration^ according to a sigmoid-like function 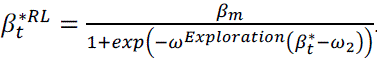. The ω_2_ parameter was fixed to 0.5. When 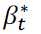 is positive, the denominator is larger than one, decreasing β*^RL^, (with lower β*^RL^ values causing more exploratory choices) whereas when it is negative the exponential is much smaller than one, hence β*^RL^ is large (causing more exploitative choices). Taken together, successive errors lead to an increase in 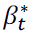, it thus more or less counts the errors, which leads via the sigmoid transformation to lower β*^RL^ hence more exploration. Conversely, successive correct choices decrease 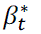, which via the sigmoid transformation reduces β*^RL^ yielding more exploitation.

*Combining SF, AE and WM mechanisms using a strategy weight.* Combining the RL mechanisms of selective forgetting and adaptive exploration with the WM component was achieved with a trial-by-trial varying strategy weight which adjusted the relative weight of the WM choice probabilities relative to the RL choice probabilities for a final choice. Previous work has shown that learning is better predicted when both, the RL and the WM component, influence choosing objects during learning ^23^. The relative influence of WM on choosing an object is determined by a weighting factor 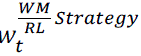 which combines the WM and RL choice probabilities into an integrated choice probability 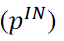. calculated as 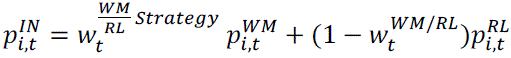. A higher 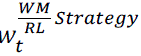 corresponds to a more prominent role of the WM component in determining the choice. We refer to 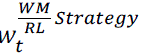 as learning *strategy weight*. The model increases the WM influence (by increasing 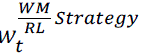) when it has sufficient capacity to hold the value of objects and when its choice probability is high relative to the choice probability of the RL component, which occurred transiently during the initial learning after a block switch when the expected values are only slowly adjusting while the working memory object value is updated immediately after a correct choice (**Figure 6K,L**).

### Selection of behavioral model

To determine whether the AE, SF, and WM mechanisms contributed to predicting task performance of the monkeys we fit models with all combinations of these mechanisms to the choices of each of the subjects of the stimulation experiment and calculated the negative log-likelihood and the Akaike Information Criterion (AIC). We fit the model to choices of the sham and the stimulation conditions but separately for each monkey and each brain areas in order to derive the best-fitting model for each monkey and area but across all experimental conditions (**Figure 5B**). The fitting minimized the Negative Log Likelihood (NLL) using the MATLAB (The Mathworks, Inc.) function *fminsearch* followed by calls to *fmincon*.

After determining that the *AE-SF-WM model*, best described the overall data, we separately fitted the *AE-SF-WM model* to the sham and the stimulation condition for the 3D task performance of each monkey. To estimate the variability of the model parameters we used a bootstrap approach that fit the behavioral data of each condition with random subsets of 80% of the blocks from that condition, generated the sequence of choice with the bootstrapped model parameter values, and extracted the static and the trial-by-trial varying parameter values of from the model generated data. We visualize the mean and standard error of the mean of these parameters in **Figure 5C-H** and **Figure 6-L**.

## Acknowledgments

This work was supported by the National Institute of Mental Health (R01MH123687). The funders had no role in study design, data collection and analysis, the decision to publish, or the preparation of this manuscript.

## Author Contributions

R.L.T., K.B.B and T.W. prepared the experiment and reconstructed the anatomical stimulation locations. R.L.T. and A.N. performed the stimulation experiments. R.L.T., C.G. and A.N. performed the electrophysiological experiments. C.G. analyzed the electrophysiological signals. R.L.T., T.W. and P.T. performed analysis, modeling and visualization. T.W. conceived the research and supervised the study. All authors contributed writing the paper.

## Data and code accessibility

Data and custom programming code for analysis and modeling is available upon reasonable request.

